# Pangenome graph analysis reveals extensive effector copy-number variation in spinach downy mildew

**DOI:** 10.1101/2024.05.30.596583

**Authors:** Petros Skiadas, Sofía Riera Vidal, Joris Dommisse, Melanie N. Mendel, Joyce Elberse, Guido Van den Ackerveken, Ronnie de Jonge, Michael F. Seidl

**Author notes:** corresponding author: Padualaan 8, Room N611, 3584 CH Utrecht, the Netherlands.

## Abstract

Plant pathogens adapt at speeds that challenge contemporary disease management strategies like the deployment of disease resistance genes. The strong evolutionary pressure to adapt, shapes pathogens’ genomes, and comparative genomics has been instrumental in characterizing this process. With the aim to capture genomic variation at high resolution and study the processes contributing to adaptation, we here leverage and expand on an innovative, multi-genome method to construct, annotate, and analyse the first pangenome graph of an oomycete plant pathogen. We generated telomere-to-telomere genome assemblies of six genetically diverse isolates of the oomycete pathogen *Peronospora effusa*, the economically most important disease in cultivated spinach worldwide. The pangenome graph demonstrates that *P. effusa* genomes are highly conserved, both in chromosomal structure and gene content, and revealed the continued activity of transposable elements which are directly responsible for 80% of the observed variation between the isolates. While most genes are generally conserved, pathogenicity related genes are highly variable between the isolates. Most of the variation is found in large gene clusters resulting from extensive copy-number expansion. Pangenome graph-based discovery can thus be effectively used to capture genomic variation at exceptional resolution, thereby providing a framework to study the biology and evolution of plant pathogens.

**Author Summary:** Plant pathogens are known to evolve rapidly and overcome disease resistance of newly introduced crop varieties. This swift adaptation is visible in the genomes of these pathogens, which can be highly variable. Such genomic variation cannot be captured with contemporary comparative genomic methods that rely on a single reference genome or focus solely on protein coding genes. To overcome these limitations and compare multiple genomes in a robust and scalable method, we constructed the first pangenome graph for an oomycete filamentous plant pathogen with six telomere-to-telomere genome assemblies of *Peronospora effusa*. This high-resolution pangenomic framework enabled detailed comparisons of the genomes at any level, from the nucleotide to the chromosome, and for any subset of protein-coding genes or transposable elements, to discover novel biology and potential mechanisms for the rapid evolution of this devastating pathogen.

## Introduction

Filamentous plant pathogens, such as fungi and oomycetes, cause devastating diseases on crop plants, resulting in severe damage in agriculture and natural ecosystems worldwide (McMullan *et al*., 2015; Hartmann *et al*., 2017; Westerhoven *et al*., 2024). Disease management strategies mainly depend on chemical pesticides and host resistances (Bourguet *et al*., 2016; Fayyaz *et al*., 2023; Zaccaron and Stergiopoulos, 2024). However, filamentous plant pathogens can rapidly overcome plant resistances and develop pesticide tolerances (Bourguet *et al*., 2016; Corkley, Fraaije and Hawkins, 2022), especially since resistant crop cultivars have often been deployed in large monocultures (Miller *et al*., 2020). These agricultural practices exert strong selective pressure on pathogens, leading to rapid diversification of pathogen populations (Möller and Stukenbrock, 2017; Mohd-Assaad, McDonald and Croll, 2019).

To establish a successful infection, pathogens need to create an environment that supports host colonization, for instance by circumventing or suppressing host immune responses (Rovenich, Boshoven and Thomma, 2014; Hartmann *et al*., 2017). To this end, pathogens secrete so-called ‘effector’ proteins that often play roles in deregulating the host immune system or in manipulating host physiology to release nutrients (Judelson and Ah-Fong, 2019). Effectors can, however, also be recognized by plant immune receptors (Cook, Mesarich and Thomma, 2015). Effector recognition often triggers hypersensitive defence responses, a process called effector-triggered immunity (Sharpee and Dean, 2016). In turn, this strong selection pressure imposed by the host immune system drives the emergence of novel or the diversification of existing effector repertoires to re-establish successful host colonization (Hartmann *et al*., 2017), leading to ongoing co-evolutionary arms races between pathogens and their hosts (Möller and Stukenbrock, 2017).

Co-evolution between pathogens and their hosts shapes pathogens’ genomes, and genomic research over the last decades has started to uncover genomic signatures of molecular processes that fuel rapid effector diversification (Ye *et al*., 2021). Comparative genomics of the oomycete plant pathogen *Phytophthora infestans* and related sister species have revealed a genome organization characterized by gene-dense genomic compartments that are largely conserved and by gene-sparse genomic compartments that are rich in transposable elements (TEs) and contain effectors and other virulence-related genes (Haas *et al*., 2009). These gene-sparse compartments are often highly variable between species and strains of the same species with an overabundance of structural variation as well as effectors and virulence-related genes that often evolve under positive selection (Raffaele *et al*., 2010). While this structure is not universal to all filamentous plant pathogens, it has been widely observed in many important plant pathogens (Dong, Raffaele and Kamoun, 2015; Thomma *et al*., 2016; Ingram *et al*., 2020; Knaus *et al*., 2020; Torres *et al*., 2020; Westerhoven *et al*., 2024). This led to the emergence of the two-speed genome model, in which genome organization is thought to facilitate the rapid diversification of virulence-related genes (Raffaele and Kamoun, 2012; Dong, Raffaele and Kamoun, 2015; Frantzeskakis, Kusch and Panstruga, 2019; Torres *et al*., 2020), thereby enabling pathogens to quickly adapt to new or changing environments and to overcome host defences. However, this model does not offer a universal explanation for the location of all effector genes, and it is yet unclear how this genome structure has evolved (Torres *et al*., 2020).

Thus far, research into the genomic diversity of specific filamentous plant pathogens has often focused on the direct comparison of genomes from multiple strains of a single species to a common, single reference genome (Everhart, Gambhir and Stam, 2021). However, comparisons to single-reference genomes can only capture a small fraction of the overall species-wide diversity, especially when genomes vary in chromosomal organization as well as gene and repeat content (Della Coletta *et al*., 2021; Garcia *et al*., 2024). The fraction of the genome that is common to all strains within a species has been termed the ‘core’ genome and the variable fraction is often referred to the ‘accessory’ genome (McCarthy and Fitzpatrick, 2019; Badet and Croll, 2020). Filamentous plant pathogens often show extensive presence/absence variation between strains, leading to considerable diversity in their accessory regions and thus potential functions (Badet and Croll, 2020). Importantly, effectors and other virulence-related genes are often enriched in the accessory genome (Badet and Croll, 2020), and single reference approaches therefore hamper the identification and subsequent characterization of pathogen effector repertoires.

To overcome potential biases introduced by single reference genomes and to better represent and study the accessory genomes of filamentous plant pathogens, genomes from multiple strains can be jointly analysed to create a pangenome (Garcia *et al*., 2024). A pangenome can be defined as the total number of unique genes discovered among genomes of the same species (McInerney, McNally and O’Connell, 2017; Badet *et al*., 2020; Hickey *et al*., 2023). However, this definition ignores the contribution of TEs and other non-coding sequences to genome diversity and adaptation (Fedoroff, 2012; Faino *et al*., 2016; Badet and Croll, 2020; Torres, Thomma and Seidl, 2021), which is particularly relevant for TE-rich accessory regions in filamentous plant pathogens (Fletcher and Michelmore, 2023). To include these regions, a pangenome needs to be defined as the total genome content of a species, thus creating a sequence-resolved pangenome. This pangenome can be represented by a variation graph where each node in the graph corresponds to a nucleotide sequence and edges between nodes indicate the direction and therefore the ‘path’ of sequences through the graph (Hickey *et al*., 2023). Consequently, genetic variation such as single nucleotide and structural variation between strains appear as ‘bubbles’ of alternative paths through the graph (Hickey *et al*., 2023).

The oomycete *Peronospora effusa* causes downy mildew, the economically most important disease of cultivated spinach worldwide (Lyon *et al*., 2016; Ribera *et al*., 2020). This pathogen has been traditionally managed by the extensive deployment of genetic disease resistances (Koike, Smith and Schulbach, 1992; Kandel *et al*., 2020). However, *P. effusa* rapidly breaks resistances of newly introduced varieties (Ribera *et al*., 2020). To date, 19 *P. effusa* races have been denominated based on their capacity to break spinach resistances (Lyon *et al*., 2016; Feng, Saito, *et al*., 2018; Klein *et al*., 2020). *P. effusa* can reproduce both sexually and asexually and thus new races can emerge both from sexual and asexual recombination (Feng *et al*., 2020; Skiadas *et al*., 2022). *P. effusa* is one of few oomycetes with a publicly available chromosome-level reference genome (Fletcher *et al*., 2022; Matson *et al*., 2022; Fletcher and Michelmore, 2023). The *P. effusa* genome is relatively small compared with other oomycetes and organised in 17 chromosomes (Fletcher *et al*., 2022). The number of chromosomes and their overall collinearity is largely conserved with closely related oomycetes, pointing to a conserved genome organisation in Peronosporales (Fletcher *et al*., 2022, 2023). The *P. effusa* genome is predicted to be composed of about 53.7% TEs and around 9,000 protein-coding genes. Of these genes, 300 are predicted to encode for secreted effectors that are enriched in TE-rich genomic regions, thus *P. effusa* seems to follow the two-speed genome model (Fletcher *et al*., 2022). Two main families of cytoplasmic effectors have been characterized in oomycetes thus far, the RXLR and Crinkler (or CRN) effector families (Haas *et al*., 2009; Saraiva *et al*., 2022). These families of effectors are characterized by the presence of conserved motifs at the N-terminus downstream of the signal peptide, which has been hypothesized to contribute to effector translocation into the host cell (Whisson *et al*., 2007; Ingram *et al*., 2020; Wood *et al*., 2020). The C-termini regions of these effectors varies significantly, and it is responsible for the effector function in the plant cell (Dou *et al*., 2008; Jiang and Tyler, 2011; Wood *et al*., 2020). Diversification of effector gene repertoires most likely drives the evolution of *P. effusa* and the rapid breakdown of host resistances (van Kogelenberg *et al*., 2015). However, comparison of the recently published chromosome-level genome assembly of *P. effusa* strain UA202013 with closely related and likely clonally evolving isolates from races 12, 13, and 14 reported limited genomic diversity (Fletcher *et al*., 2021). To date, however, chromosome-level genome assemblies of multiple *P. effusa* strains are still lacking and thus the evolutionary processes that might contribute to emergence of novel races are largely unknown.

Here, we leveraged and expended on a pangenome graph-based approach to enable comprehensive, reference-free analysis of chromosome-level genome assemblies of multiple genetically diverse *P. effusa* isolates. We show that the chromosome structure and most protein-coding genes are largely conserved. In contrast, most genomic variation is caused by TEs, but also effector and other virulence-related gene repertoires are highly dynamic. These genes are often located in clusters of highly similar gene copies, which reveals their recent copy-number changes and is the leading cause for their variation between the isolates. Our pangenome approach provides a framework to compare isolates of microbial pathogens accurately and efficiently, which will be essential to understand how these are able to rapidly break host resistances in the future.

## Results

### Chromosome-level genome assemblies of six *Peronospora effusa* isolates reveal a highly conserved genome structure

To fully capture the genomic variation of spinach downy mildew *P. effusa* by constructing a pangenome graph, we first sought to create gapless chromosome-level genome assemblies of a suite of diverse *P. effusa* isolates by a combination of Nanopore long-reads and short Hi-C paired-end reads. We selected *P. effusa* isolates that belong to six denominated races with the aim to capture large parts of the genomic variation present between all thus far sequenced *P. effusa* isolates (Skiadas *et al*., 2022). We selected one isolate for each of the three phylogenetic clusters we have previously identified (isolates classified as *Pe1*, *Pe5*, and *Pe14*), and three that most likely emerged via sexual reproduction between isolates of those phylogenetic clusters (isolates classified as *Pe4*, *Pe11*, and *Pe16*) (Skiadas *et al*., 2022) (S. Figure 1).

For all six isolates, we used Nanopore, Hi-C, and Illumina sequencing data to create gapless assemblies with 17 chromosomes, either with or without an additional 18^th^ chromosome, with overall genome sizes ranging between 57.8 and 60.5 Mb (Figure 1A, S. Figure 2, S. Figure 3, S. Table 1, S. Table 2, S. Note 1). The 17 assembled chromosomes have telomeric repeats with the repeat motif ‘TTTAGGG’ on both ends, suggesting that we successfully obtained complete and chromosome-level genome assemblies for all six isolates. The assembly completeness, evaluated by Benchmarking Universal Single-Copy Ortholog (BUSCO) analysis using the Stramenopiles database (v. odb10), revealed a 99% BUSCO completeness score, indicating that we successfully captured the protein-coding regions.

Hi-C data provides chromatin contact information, based on which, one can extract details about the 3D organisation of chromosomes in the nucleus (Torres *et al*., 2023). The Hi-C heatmaps of *P. effusa* show increased interaction frequencies between all chromosomes with a characteristic pattern that matches the Rabl chromosome configuration (Hoencamp *et al*., 2021a; Torres *et al*., 2023a); all the telomeres co-localize within the nucleus, shown as dots in the heatmap, and similarly all centromeres co-localize, indicated by the characteristic ‘x’ pattern in the Hi-C heatmap (Figure 1C, S. Figure 4). The Rabl configuration has thus far only been found in species without the Condensin II complex, which organises the nucleus into chromosomal territories (Hoencamp *et al*., 2021a; Torres *et al*., 2023a), and based on sequence similarity searches, Condensin II subunits seem to be absent in *P. effusa*. By using the centromeric interactions visible in the Hi-C heatmap, we identified the approximate locations of centromeres for each chromosome (S. Table 2). These centromeric interactions are approximately 250 to 300 kb long, which is in line with centromeric regions (211 to 356 kb in size) measured in *Phytophthora sojae* (Fang *et al*., 2020). Importantly, centromeric regions, except for chromosomes 1 and 15, are enriched for copies of a *Copia*-like LTR transposon (Figure 1A), which has been previously identified to occur at centromeres in different oomycetes (Fang *et al*., 2020).

The previously published genome assembly of the *P. effusa* isolate UA-202013 has 17 chromosomes (Fletcher *et al*., 2022), and is comparable in total genome size as well as gene and repeat content to the here assembled *P. effusa* isolates. Since the overall chromosome organization of oomycetes is thought to be highly conserved (Fletcher and Michelmore, 2023), we sought to investigate the co-linearity between our *Pe1* isolate, the UA-202013 isolate, and chromosome-level genome assemblies from *Bremia lactucae*, *Peronosclerospora sorghi*, and *Phytophthora infestans* (Fletcher *et al*., 2019, 2022, 2023; Matson *et al*., 2022). To this end, we performed whole-genome alignments of these genomes based on sequence similarity and on relative position of protein-coding genes (S. Figure 5A). The *Pe1* and UA-202013 *P. effusa* isolates are completely collinear, and between species the chromosome structure remains largely conserved apart from few chromosome fusions and inversions of large chromosomal regions. We similarly observed that our six *P. effusa* isolates are nearly completely co-linear (S. Figure 5B). Thus, the chromosomal organization between different oomycetes and especially between isolates of the same species are highly conserved (Judelson, 2014). We nevertheless also observed two large structural rearrangements (inversions), one in *Pe5* at the beginning of chromosome 17 and one in *Pe11* at the end of chromosome 1 (S. Figure 5B), indicating that chromosomal rearrangements have nevertheless occured.

**Figure 1.**
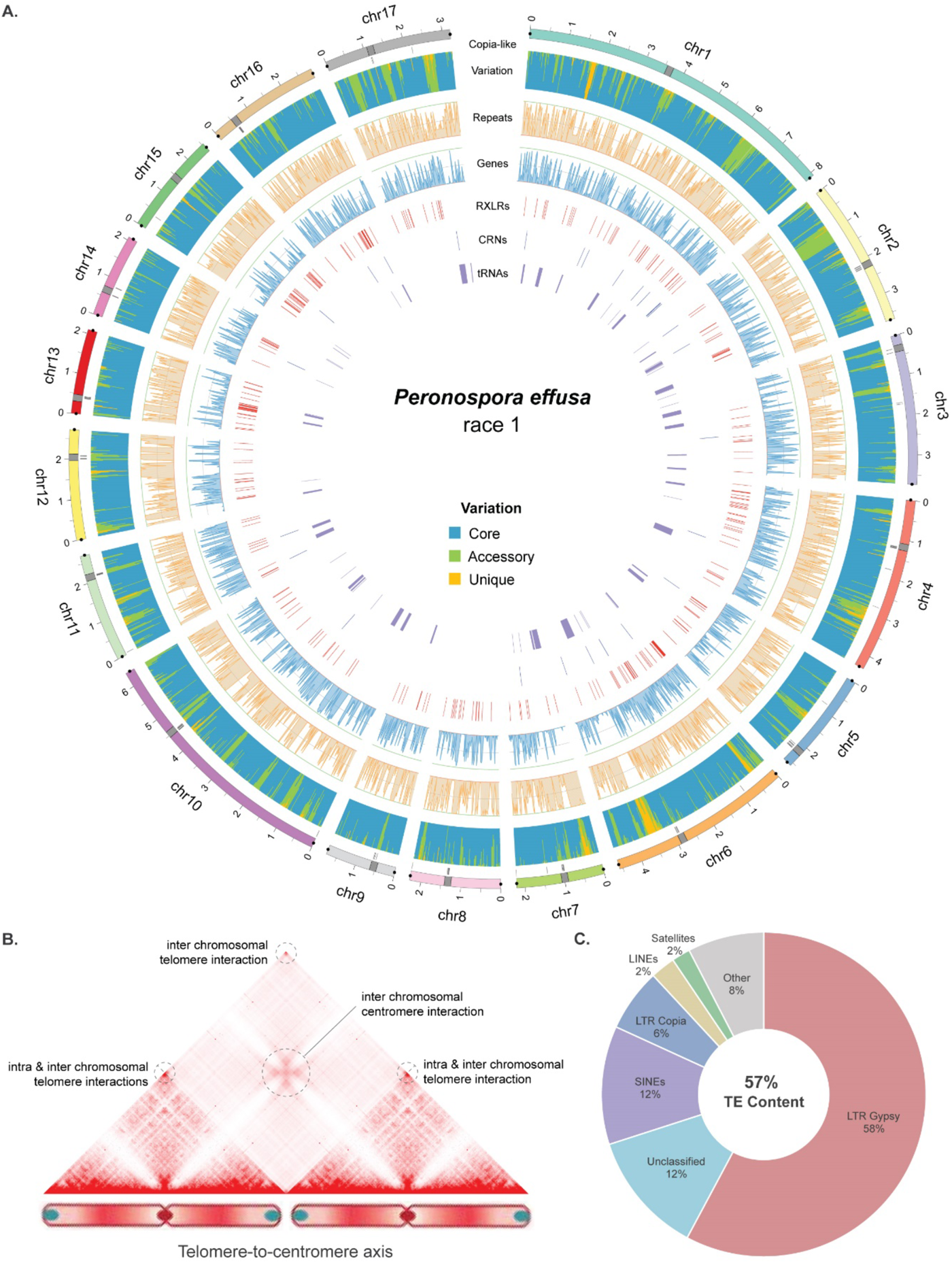
Chromosome-level genome assembly of *Peronospora effusa* race 1 (*Pe1*). **A**. The genome assembly of *Peronospora effusa* race 1 (*Pe1*) is visualised in a circular plot, individual tracks, starting from the outside to the inside: i) 17 chromosomes are shown in different colours, a grey rectangle point the location of the centromeres, grey lines underneath each chromosome show the location of the centromere-specific *Copia*-like element, which has been previously observed to be enriched in centromeres (Fang *et al*., 2020), and dots at either side of the chromosomes highlight the presence of telomeric repeats. ii) Stacked bar plot shows the genetic variation in the six *P. effusa* isolates, with blue representing conserved regions, green regions that are present in at least two isolates, and orange unique regions present only in *Pe1*. iii) Line graph shows the coverage of repeat (orange) and gene (blue) content summarized in non-overlapping 20 kb windows. iv) Lines indicate the position of RXLR (red), CRN (green), and tRNA (purple) genes. **B.** Hi-C heatmap displays the spatial interactions between chromosomes in the nucleus. Genome-wide Hi-C data is aggregated over two chromosomes to highlight inter- and intra-chromosomal telomeric interactions and inter-chromosomal centromeric interactions. This interaction pattern suggests that *P. effusa* chromosomes are organized within the nucleus in a Rabl chromosome configuration (Hoencamp *et al*., 2021b; Torres *et al*., 2023b) (S. Figure 3). **C.** Transposable element (TE) content in the *Pe1* genome assembly is shown separated over different TE families as percentage of the total length of TEs annotated.

### Pangenome graph-based comparison of six *Peronospora effusa* genomes reveals overall high conservation but highly variable TE and effector genes

To enable the comparisons between the *P. effusa* chromosome-level genome assemblies and to limit possible reference biases introduced by comparisons to a single reference genome, we created pangenome graphs based on progressive Cactus (Hickey *et al*., 2023). The conserved structure of the chromosomes in *P. effusa* (S. Figure 5B) enabled us to generate a separate pangenome graph for each chromosome. Collectively, the 17 pangenome graphs have 1,948,729 nodes and 2,635,318 edges, which is only a small fraction of all theoretically possible connections between the nodes, indicating that these graphs are simple in structure. The total size of the graph is 76.6 Mb, 32% larger than the average *P. effusa* genome assembly (58 Mb), suggesting that while most of the genome is conserved, there are significant strain-specific genomic regions (Figure 2A).

To generate a homogenous framework to study the genetic differences between the six *P. effusa* isolates, we created a common TE library, and we exploited the pangenome graph to perform a joined structural annotation for the protein-coding genes on the six genome assemblies (S. Note 2). These methods ensure that the TE annotations are based on shared TE families and that genes that are present in identical regions in the pangenome graph will be annotated consistently between the isolates (e.g. identical intron-exon structure and ORF position) (Figure 2B steps 1-3). This resulted in the annotation of 9,916 to 10,364 protein-coding genes (on average 67% being functionally annotated with interproscan v. 5.52-86.0) for each isolate, adding between 18 to 459 genes that were missed by the individual gene annotation of each isolate. These genes were then assigned into groups based on their position on the pangenome graph, thus creating 12,739 single-copy gene orthogroups (Figure 2B step 4, S. Note 2). To identify effector candidates, we searched the predicted protein-coding genes and additional open reading frames for those encoding proteins with a predicted signal peptide and for RXLR and CRN amino acid motifs. We identified 351 to 443 putative RXLR and 38 to 68 putative CRN effectors per isolate (Figure 1A, S. Table 2). To study the distribution of these genes in the genome, we searched for genes that share the same functional annotation and are physically clustered together in the genome. Of the 1,393 functional groups with at least two genes in *Pe1*, 155 occur in at least one cluster. Additionally, of the 59 functional groups with at least 20 genes, 13 have at least 15% of their genes in a cluster. These observations suggest that physical clustering of functionally related genes is a common phenomenon.

To uncover the genomic variation between the six *P. effusa* isolates, we parsed the 17 pangenome graphs and inferred that 60% (around 45.6 Mb) of the *P. effusa* pangenome is conserved, 24% (18.6 Mb) is found in two or more isolates, and 16% (12.3 Mb) is unique for single isolates (Figure 2A, Figure 3A). Most of the observed variation originates from variants longer than 50 bp (87.7% of total variation size, 27.1 Mb), but single nucleotide variants (SNVs) are most frequent (76.5% of total variation number, 1.15 million) (Figure 3A). As expected, protein-coding regions are generally highly conserved (89.0%) compared with regions annotated as TEs (48.7%). This variation in TEs in mostly caused by large sequence variants (>1 bp) that are more abundant in TE regions (42%, 277,151) than in protein-coding regions (5%, 31,786), suggesting that large variation is most often occurs in TEs or directly caused by TE activity.

Genetic variation in the pangenome graph can be specific to only a single isolate or shared between multiple isolates. We observed that 20.2% (329,387) of the variation in TE regions is unique to only one isolate, while only 12.6% (30,948) of the variation in protein-coding regions is unique, suggesting that the addition of more diverse *P. effusa* isolates would differentially impact the number of observed genes and transposons. We therefore sought to visualize the variation discovered on the pangenome graph as a saturation plot for the whole pangenome (Figure 3B). The pangenome is clearly open (not saturated) for TEs and explains 80% of the observed variation between the *P. effusa* isolates. The TE content in *P. effusa* is highly dynamic and shaped by continuous TE expansions and deletions. To investigate the timeline of repeat expansion, we calculated the divergence of individual TE copies to their TE consensus sequence (Kimura distance), which uncovered a recent expansion of LTR *Gypsy* and *Copia* elements, as well as an older expansion of LTR *Gypsy* elements (Figure 3C, S. Figure 6). The largest LTR *Gypsy* family has around 250 copies per isolate and is found in all chromosomes. These copies have a Kimura distance of only 0.002 and 60% occur within accessory or unique regions, suggesting that this TE family remained highly active after the divergence of the different *P. effusa* isolates. In contrast to the open TE pangenome, the pangenome for genes is nearly closed, demonstrating that by analysing only six diverse *P. effusa* isolates we have successfully captured most protein-coding genes in the population (Figure 3B). Effector genes though are much more variable than the rest of the genes, due to extensive copy-number variation. Since we have defined single copy orthogroups, copy-number variation is accounted for in saturation plots, revealing an open pangenome for effector genes (S. Note 2) (Figure 3B).

Chromosomes do not only differ in size, but also in the proportion of conserved regions, as well as repeat and gene content. We used these characteristics to study individual chromosomes with principal component analyses (PCA), which revealed four distinct groups of chromosomes (Figure 3D). The first group consists of highly conserved chromosomes (69 - 87% core), which includes the most repeat poor (52 - 62%) and smallest (1.7 - 2.3 Mb) chromosomes; chromosome 4 is an outlier based on chromosome size (4.3 Mb) and repeat content (70%) because it contains an older repeat expansion that affected its size but is shared between all isolates (S. Figure 7; Panel Pe5_Chr4). The second group is characterized by a high repeat content (68 - 70%). The third group consists of the least conserved chromosomes (45 - 60% core), with some of the largest (3.2 - 4.6 Mb) chromosomes. Lastly, the fourth group consists of the two largest chromosomes 1 (8.1 Mb) and 10 (6.3 Mb), which clearly grouped apart of the other chromosomes even though these chromosomes are similarly conserved to most chromosomes, and they have an average repeat content (chromosome 1: 54% core, 61% repeats; chromosome 10: 58% core, 59% repeats). Interestingly, while most *P. effusa* chromosomes are completely syntenic with the chromosomes of *Bremia lactucae*, this is not the case for chromosome 10 (Fletcher *et al*., 2022), which suggests that chromosome 10 and most likely also chromosome 1 might be the result of a fusion of two smaller ancestry chromosomes, similar to other chromosome fusion events that have been observed in Peronosporaceae (Fletcher *et al*., 2022, 2023) (S. Figure 5A).

**Figure 2.**
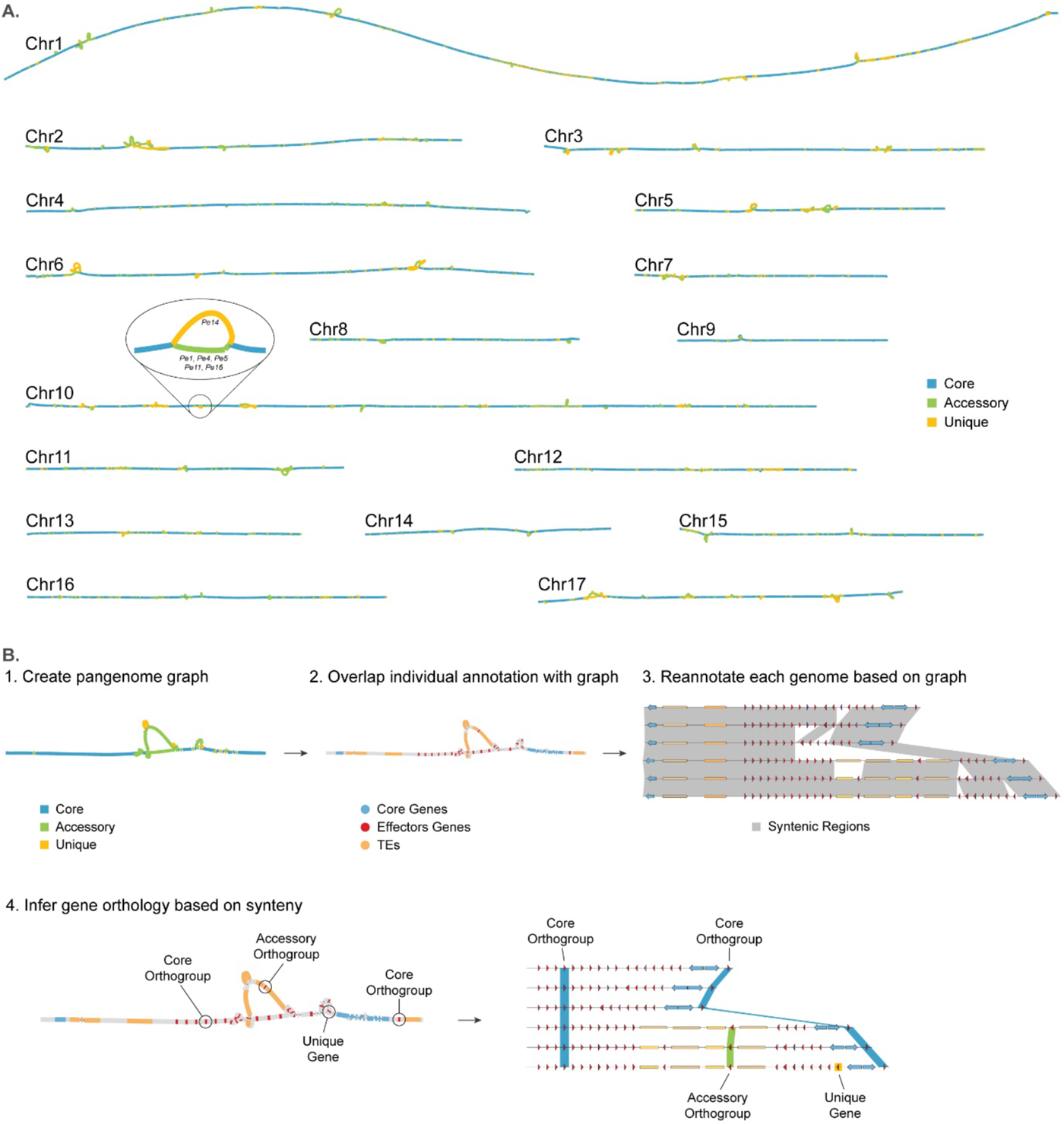
Reference-free genome annotation and whole-genome comparisons of six *Peronospora effusa* isolates with pangenome graphs. **A.** Pangenome graphs of the 17 chromosomes present in all *P. effusa* isolates. As an example of the graph structure, a variable region of chromosome 10 is highlighted, indicating the corresponding isolates for each accessory (green) and unique (orange) region in the graph. **B.** Step-by-step description of the pangenome graph analysis to reannotate and compare all protein-coding genes, including effector candidates. 1. The genome assemblies are used to create and annotate (determine the variation of each region) the pangenome graph; 2. Gene annotations are projected onto the pangenome graph; 3. All genome assemblies are re-annotated based on their alignment in the graph and associated gene annotation; 4. The overlap of genes on the graph is used to create orthologous gene groups based on synteny.

**Figure 3.**
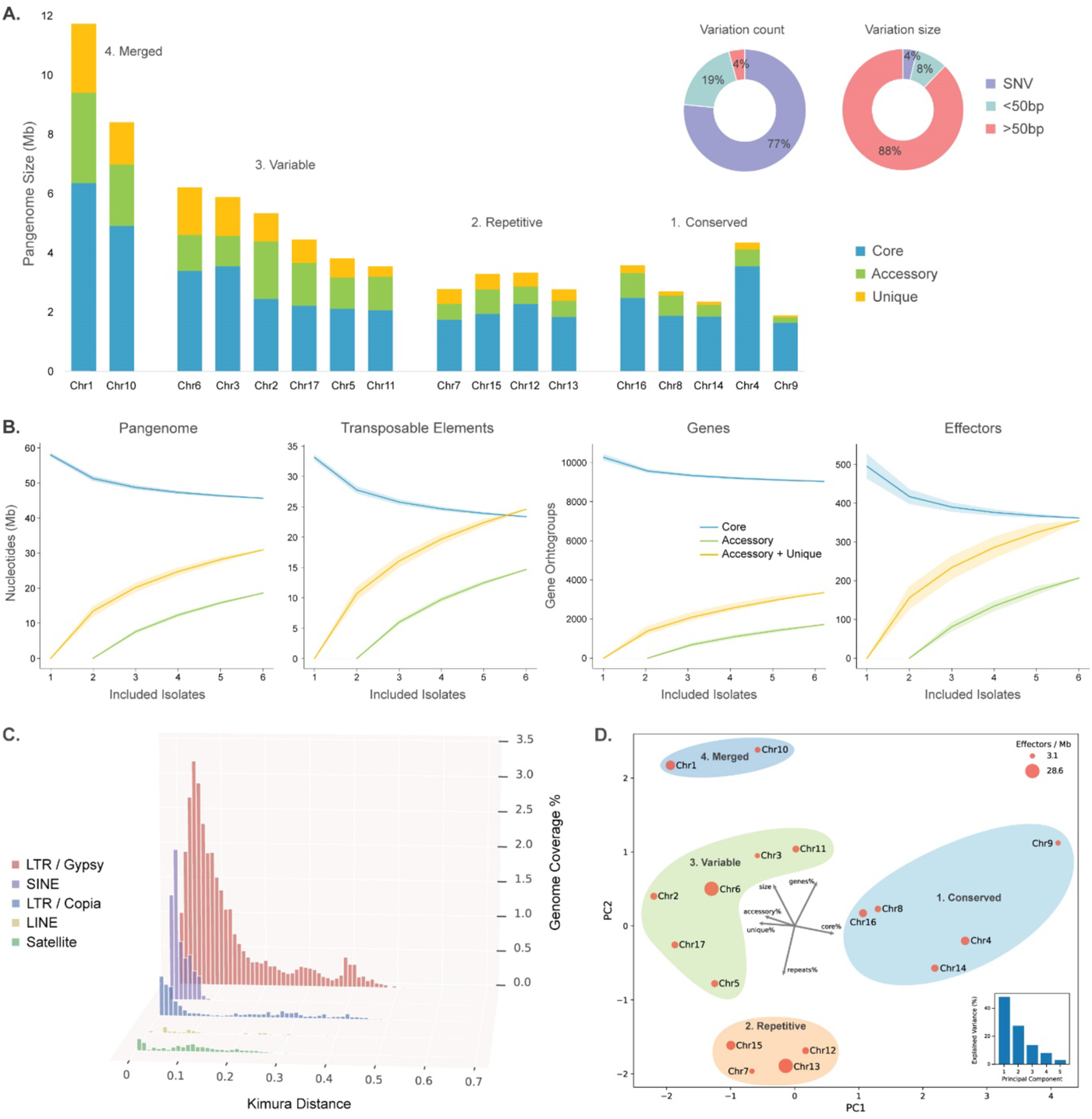
Pangenome graph based comparison of six *Peronospora effusa* isolates. **A.** The 17 chromosomes of *P. effusa* pangenome variation size represented in a stacked bar plot. The variation observed in the pangenome graph analysis for count and total size represented in two pie charts. **B.** Saturation plots based on the pangenome graph at the nucleotide level for the whole genome and transposable elements, and at the gene level for all genes and effector candidates. **C.** The transposable element landscape based on Kimura distances for the five largest TE superfamilies for *Pe1* are shown in order of coverage of the genome. Kimura distance is the measure of divergence between individual TE copies and the corresponding TE consensus sequence (Kimura, 1980), i.e., the lower the Kimura distance, the more similar the copy is to the consensus and thus the more recently it was most likely copied. **D.** Principal component analyses of the 17 *P. effusa* chromosomes based on their percentage of genes, repeats, core, accessory, and unique region and their total size. The size of each point corresponds to the number of effector genes per Mb on each chromosome. The chromosomes are clustered in four distinct groups.

### *Peronospora effusa* has a variable and highly repetitive accessory chromosome

The genome assembly of *P. effusa* isolate UA-202013 as well as the here assembled isolates *Pe1* and *Pe14* have 17 chromosomes (Figure 1) (Fletcher *et al*., 2022). However, we also assembled an additional complete chromosome in *P. effusa* isolate *Pe5*, named chromosome 18. This chromosome is similar to other chromosomes as it is 2.1 Mb long, has full diploid coverage, has a centromere based on the Hi-C data, and is flanked by telomeric repeats (S. Figure 3). In contrast to other chromosomes, however, it is mostly composed of TEs of the LINE and LTR superfamily (87% TE content compared with 54% genome-wide average), has higher GC content (54% compared with an average of 48%), and the two chromosomal arms are the reverse complement of each other (Figure 4A). Moreover, unlike other chromosomes, this one shares almost no sequence similarity with any other chromosome, since all the annotated LINE, satellite, and unknown repeats are unique to chromosome 18. Only four significant matches were found with other chromosomes, corresponding to copies of two LTR-*Gypsy* families, which had a recent expansion and are among the most abundant TE families in *P. effusa*. Near the telomeres where the two LINE/L1 clusters are located, we annotated 18 putative protein-coding genes, nine in each of the chromosome arms. These sequences lack similarity to any other predicted proteins of *P. effusa* or any other known proteins from public databases. Since these are also not expressed, we concluded that these are most likely non-functional. Notably, we also assembled additional contigs in *Pe4*, *Pe11*, and *Pe16* that share sequence similarity with chromosome 18 of *Pe5*. However, these are 30-36% shorter, have only a single telomeric repeat, and miss half of a chromosomal arm that is unique for *Pe5* as shown in the pangenome graph (Figure 4B). We similarly observed protein-coding genes annotated in each of these contigs, yet we could not identify sequence similarity between different isolates, further corroborating that these predicted protein-coding genes are likely non-functional.

To better understand the occurrence and possible origin of this accessory chromosome, we used publicly available short-read data from whole-genome resequencing experiments for in total 32 *P. effusa* isolates (Feng, Lamour, *et al*., 2018; Fletcher *et al*., 2018; Klein *et al*., 2020). Based on read mapping to the *Pe5* chromosome 18, we identified similar accessory chromosomes in nine *P. effusa* isolates, including sequencing data of three independent isolates that are classified as race 13 (*Pe13*, *R13*, and *Pfs13*) (Feng, Lamour, *et al*., 2018; Fletcher *et al*., 2018; Klein *et al*., 2020). In eight *P. effusa* isolates, such as *Pe4*, *Pe11*, and *Pe16,* chromosome 18 is partially present (40-60% of the entire chromosome is covered), and in two isolates we could find only traces of the genetic material assigned to *Pe5* chromosome 18 (8-10% coverage) (Figure 4C). Interestingly, based on the known relationship between the isolates (Skiadas *et al*., 2022) (Figure 4D), we could not identify any traces of chromosome 18 in data from isolates of clusters i and iii as well as in *Pe10*, *Pe17*, and *NL-05*, suggesting that chromosome 18 was likely present in isolates that form cluster ii and that this chromosome was subsequently lost or degraded in a subset of isolates, possibly due to recombination with isolates from cluster i and iii (Figure 4D). Our analysis shows that chromosome 18 is present in the closely related beet downy mildew *Peronospora farinosa* f. sp. *betae* (ES-15) (Figure 4C), indicating that it was present in their last common ancenstor and has been recently lost in many of the *P. effusa* isolates, or gained from cluster ii after the differentiation of these *P. effusa* isolates.

**Figure 4.**
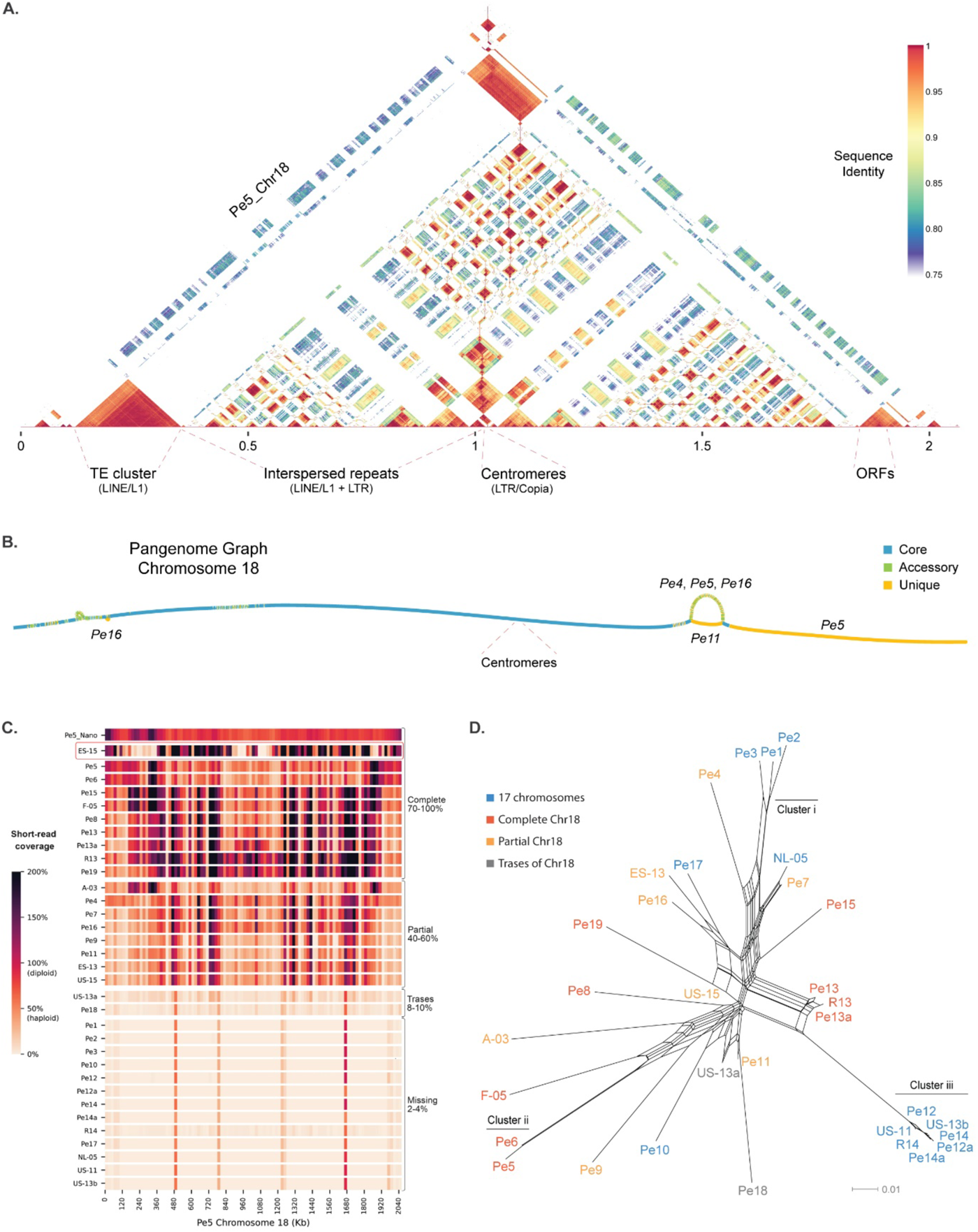
A highly repetitive chromosome is present in a subset of *Peronospora effusa* isolates. **A.** Self-alignment of *Pe5* chromosome 18 in 1.5 kb windows shows the repetitive nature of the accessory chromosome and highlights the two chromosomal arms that are reverse compliment to each other. **B.** Pangenome graph of accessory chromosome 18 in *Pe4*, *Pe5*, *Pe11*, and *Pe16*. **C.** Presence of chromosome 18 in 32 *P. effusa* isolates and the beet downy mildew (indicated with red box) based on the short-read coverage compared with the *Pe5* chromosome assembly. When the coverage is 100% of the average coverage over the whole genome, it indicates that the chromosome is diploid. The top row displays the coverage of the Nanopore long reads of *Pe5*, while the remaining rows show coverage based on Illumina short-read data for each analysed *P. effusa* isolate. **D.** The Presence of chromosome 18, as determined in C, is shown in the context of the *P. effusa* phylogeny. Similarity between *P. effusa* isolates is represented in a neighbor-net phylogenetic network; the branch lengths are proportional to the calculated number of substitutions per site.

### Extensive variation in repeat-rich regions is caused by rapid repeat expansion and contraction

The most extensive variation in the pangenome graph occurs at gene clusters formed by genes annotated as tRNA genes (Figure 1A), which appear in the pangenome graph as large ‘bubbles’ of unique and accessory regions (Figure 2A, Figure 5A). These clusters are found in multiple locations across all 17 core chromosomes (33-39 per isolate) and contain the vast majority of all tRNA genes annotated within a genome (around 99.6% of the more than 6,000 annotated in each *P. effusa* isolate). Individual clusters differ in size and can contain between ten to 970 tRNA genes. Notably, the tRNA genes of each cluster are nearly identical (>99.99% identity), while they share less similarity (<70% identity) with other tRNA genes identified outside the clusters. In between the tRNA genes, we discovered conserved regions with a highly similar open reading frames (ORFs; >98% identity), but single nucleotide deletions cause frame shifts changing the size of the predicted encoded proteins (Figure 5B). Interestingly, some of these ORFs are annotated as complete genes of around 300 nucleotides that potentially encode a protein with a signal peptide and an RXLR amino acid motif that is also found in effector candidates.

One of the biggest tRNA clusters can be found on chromosome 6, where a cluster of Arginine tRNAs was initially annotated. In *Pe1*, 570 tRNAs (72 bp in length) in intervals of 645 bp were identified in a single 412 kb region (Pe1_Chr6:3.2-3.6 Mb) (Figure 5B). The alignment of all copies of this 717 bp repeated sequence revealed that the tRNA gene is 100% identical while the remainder of the sequences is generally less conserved (94.9-98.9% identity). Within the cluster, we observed multiple subgroups of repeats (50-200 repeats). While within each of these groups individual copies are almost identical (>99% identity), indicating that these expanded recently, we observed significant differences between copies of different subgroups (94-98% identity) and these subgroups are seperated by even less conserved sequences (80-90% identity) (Figure 5B). Moreover, in *Pe1*, we identified two cases where the repeated sequence is reversed (Figure 5B ii. and iii.). These inversions along with the changes in sequence identity are indicative for multiple and separate expansions of this 717 bp sequence within this cluster, which is unique to this location in chromosome 6.

In the other five isolates, the same repeated sequence as in *Pe1* can also be found, and in the pangenome graph we observed that the first 80 kb of this region is highly conserved between all isolates. In contrast, however, the remaining region is highly variable with different expansion patterns and large differences in the size of the cluster, ranging from 412 kb in *Pe1* to only 129 kb in *Pe14*. According to the pangenome graph, isolates *Pe16*, *Pe11*, and *Pe4* follow a similar path through the graph as *Pe1*, although *Pe1* has additional unique regions (Figure 5A). These three isolates have a similar but shorter path compared with *Pe1*, including the two inversions (Figure 5B). In contrast, *Pe5* has a unique path, starting from point (v.), which is caused by an expansion that is different compared with other isolates. Similarly, *Pe4* appears as a much shorter unique path, starting from the point of inversion (ii.), because the expansion in *Pe14* is much shorter and the inversions observed in other isolates are not present. These extensive differences between the *P. effusa* isolates suggests that these repetitive elements are continuously expanding and contracting.

The repetitive tRNA sequences and the coupled ORFs that we uncovered in the *P. effusa* genomes resemble short interspersed nuclear elements (SINEs) that have been characterised in mammals, plants, and insects, but not in oomycetes (Han *et al*., 2021). Notably, the rapid and continuous expansion that we observed here resembles the activity of TEs rather than tRNAs and protein-coding genes. SINEs are Class I TEs, propagating by copy-paste mechanism, although they do not encode any proteins and thus are depedend on other TEs for their expansion (Kanhayuwa and Coutts, 2016). Most SINEs are caracterised by the presence of a tRNA-like sequence at the 5’ terminal region, by a central conserved region, and by a relatively short sequence of 200-700 bp (Kanhayuwa and Coutts, 2016). The taxonomic distribution of a specific SINE family is often clade-specific and does not expand to more distantly related taxonomic groups (Han *et al*., 2021), thus we were not able to match the sequences found in *P. effusa* with any known SINE sequences. Based on these observations, we therefore propose to classify these sequences as SINE-like. To annotate them throughout the genomes, we extracted consensus sequences from each cluster ranging from 284 to 1,070 bp in size, and added them to our *P. effusa* common TE library. The TE annotation with these sequences revealed that 5.4 to 5.9% of the genome assembly in each of the six *P. effusa* strains is composed by these SINE-like sequences, which have undergone recent expansions (Figure 3C, Supplementary Table 2), thus explaining the observed differences between the isolates. Although, SINEs have not yet been characterized in depth in oomycetes, a similarly large amount of tRNA genes was recently reported in *Phytophthora infestans* (Matson *et al*., 2022), suggesting that SINE-like elements are commonly present and particularly active in Peronosporaceae.

**Figure 5.**
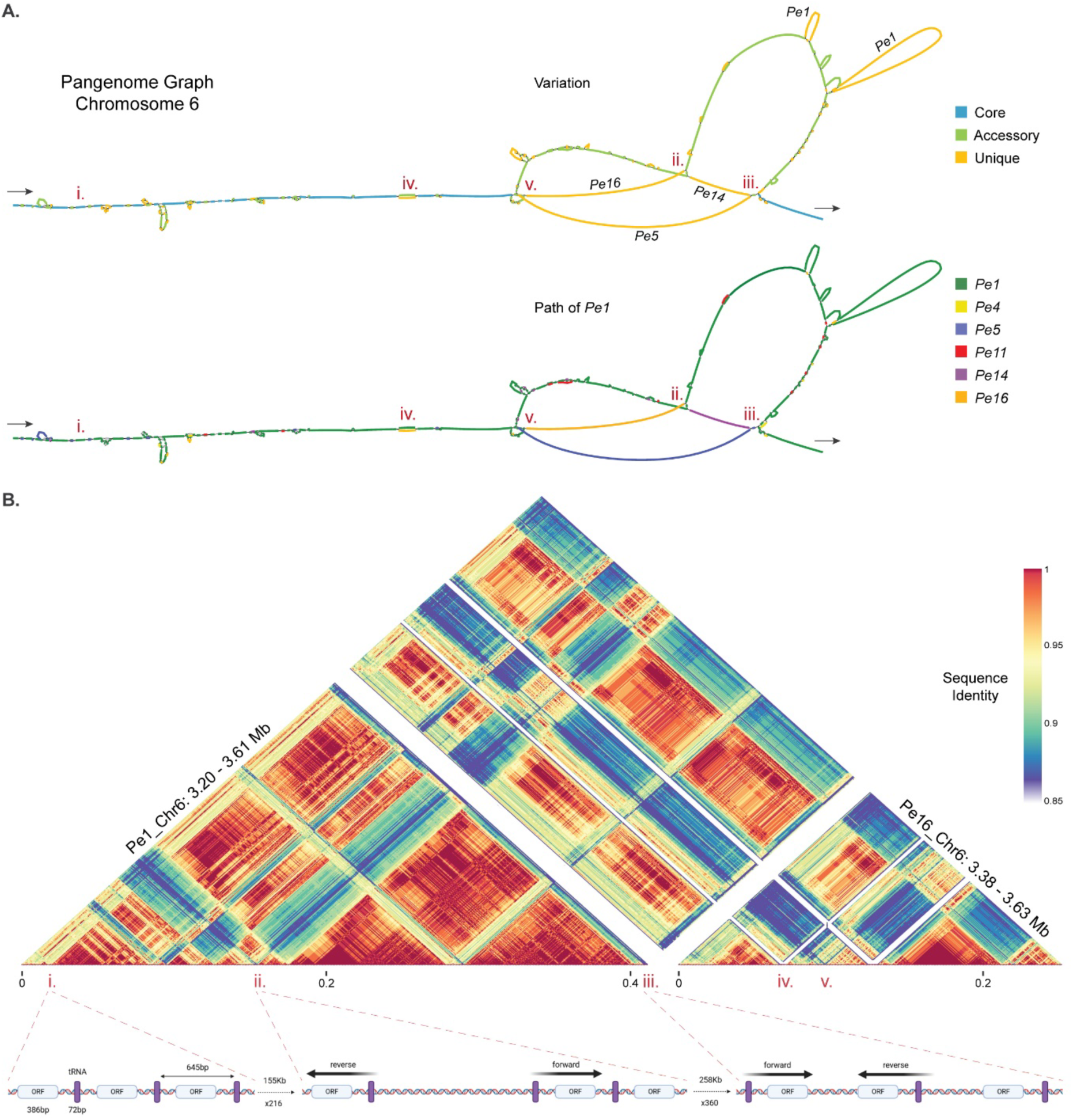
The rapid expansion of SINE-like elements causes extensive genomic variation between *P. effusa* isolates. **A.** Part of chromosome 6, 0.5 Mb in size, is visualised by two pangenome graphs, with the start and end of the graph indicated with arrows. The first graph represents the variation between the isolates and shows core (blue), accessory (green), and unique (orange) regions. In large unique regions, the name of isolate with the specific unique region is indicated. The second graph highlights the path of *Pe1* (dark green), and the unique regions of other isolates are coloured (*Pe4*: yellow, *Pe5*: blue, *Pe11*: red, *Pe14*: purple, *Pe16*: orange). Regions of interest in A and B are highlighted by roman numerals (i-v). **B.** The corresponding region of *Pe1* (Chr6: 3.20-3.61 Mb) and *Pe16* (Chr6: 3.38-3.63 Mb) is aligned in 717-bp windows, which matches the size of the identified SINE-like repetitive element. The alignment is visualised by a heatmap with the colour representing the sequence identity of each alignment from 85% (blue) to 100% (red). A schematic of the observed structure of this cluster based on the annotation of tRNA genes and ORFs is depicted below the alignment and the corresponding regions in the heatmap are indicated with red dashed lines.

### Variation in pathogenicity-related genes originate from changes in gene copy-number in gene clusters

By plotting all genes of each isolate in their chromosome position, we observed that most of the gene variation between isolates is concentrated in few, highly variable genomic regions (Figure 6A, clustering of green and yellow bars). By using the synteny-based gene orthogroups on the pangenome graph, we calculated the ratio of substitution rates at non-synonymous and synonymous sites (dN/dS). Overall, genes are highly conserved with 73% being core and 72% having dN/dS values lower than one, indicating that they evolve under negative selection (average dN/dS 0.83). In contrast, genes encoding RXLR and CRN effector candidates show extensive variation with only 50.5% found to be core and less than half (46%) evolve under negative selection and 54% show signs of positive selection (average dN/dS 1.49) (Figure 6B). Interestingly, effectors that are not part of physically co-localizing gene clusters show a distribution of dN/dS values that is comparable to the rest of the genes, and 67% evolve under negative selection (average dN/dS 0.98). In contrast, the dN/dS values of effectors that are clustered in the genome have a bimodal distribution and only 46% evolve under negative selection (average dN/dS 1.81). Thus, most of the variation observed in effectors originates from differences in gene copy-number, mainly found in distinct gene clusters (Figure 6C, S. Figure 8). For example, we observed an effector gene cluster at the beginning of chromosome 6 where *Pe5* has a unique expansion with 45 effectors, with most genes having elevated dN/dS values (average dN/dS 2.15). We also observed two less variable effector gene clusters in the middle of chromosome 13 where *Pe11* and *Pe14* share nine accessory effectors and *Pe14* has two unique effectors (average dN/dS 1.31 and 0.57) (Figure 6A).

Similar to effectors, other genes encoding proteins with functions commonly associated with pathogenicity show variation between isolates that is driven by gene copy-number changes specifically in gene clusters. Most notably, we identified 21 genes encoding necrosis-inducing proteins (62% core), which are known to induce plant cell death or suppress plant immune responses, but in oomycetes are known to be nontoxic (Cabral *et al*., 2012; Feng *et al*., 2014; Seidl and Van Den Ackerveken, 2019). Moreover, we discovered 22 genes related to glycoside hydrolase family 12 in a single cluster in the genome (45% core), known to be present in many bacteria and fungi (Zhu *et al*., 2019), and 17 papain family cysteine protease genes (76% core) that are known to be involved in virulence or defence, amongst many other functions (Ozhelvaci and Steczkiewicz, 2023). When considering genes that share the same functional annotation, those that are physically clustered in the genome are much more similar to each other than genes outside clusters (68% vs 32% protein identity). Around half of the clustered genes are variable between the isolates (52%), while most of the genes outside clusters are conserved (94%) (Figure 6C, S. Figure 8B), suggesting that most variation between the isolates originates from recent copy-number changes in gene clusters.

**Figure 6.**
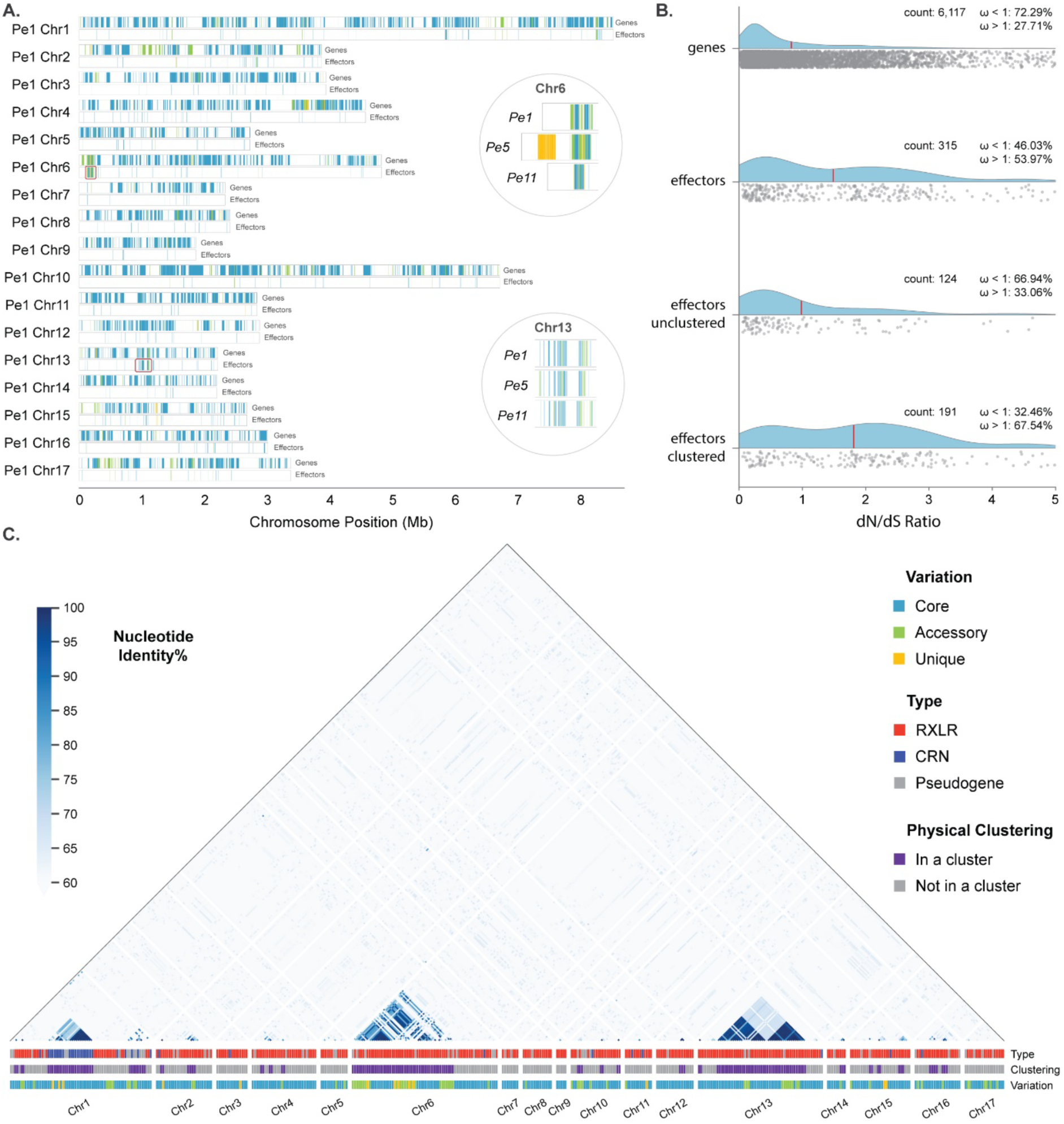
Effector variation is caused by gene copy-number changes in effector gene clusters. **A.** Gene positions in each chromosome of *Pe1* for genes and genes encoding (RXLR and CRN) effector candidates are coloured based on their conservation, blue for core, green for accessory, and orange for unique genes. Two effector clusters are highlighted on chromosome 6 and 13, and their comparisons between *Pe1*, *Pe5*, and *Pe11* is shown. **B.** Gene evolution measured for each orthogroup with dN/dS ratio (ω) shown for all genes, all effectors as well unclustered and clustered effectors. The average value is indicated by a red line. **C.** Heatmap of the nucleotide identity of the 445 annotated candidate effector genes in *Pe1* highlighting the high sequence similarity between clustered effectors. The type of the encoded effector, whether the gene belongs in a cluster, and the variation of the gene based on the pangenome graph is indicated below the heatmap.

### Changes in gene copy-number in effector clusters are associated with TE activity

The largest group of genes associated with pathogenicity, RXLR effectors, are highly variable between the isolates, which is mostly caused by gene copy-number variation within physical gene clusters. Of the total 601 RXLR orthogroups in the pangenome 47% (283) occur in clusters, but 66% (192) of the 292 variable RXLR orthogroups are found in clusters of two or more effectors. For each isolate, we observed 15 clusters with between three to five gene copies and three RXLR effector clusters with 14 or more copies, one in the beginning of chromosome 6 (Pe1_Chr6: 71-238 kb) with 30 (*Pe11*) to 111 (*Pe5*) copies, and two more at the centre of chromosome 13 (Pe1_Chr13: 908-1076 kb) that encompasses 24 (*Pe1*) to 26 (*Pe16*) copies and 14 (*Pe5*) to 23 (*Pe14*) copies, respectively. These clusters consist of almost identical sequences (>96% protein identity of genes within a cluster), indicating recent gene expansion within these clusters.

The two effector clusters in chromosome 13 are present in an otherwise highly conserved region (Figure 7A). Variation between *P. effusa* isolates is primarily caused by single gene deletions or insertions (Figure 7B, S. Figure 9). In addition to copy-number variation, gene sequences also differ due to small sequence changes, and we observed ten examples of pseudogenization caused by single nucleotide deletions and subsequent frame shifts. Moreover, all the 114 effector genes found in the second RXRL cluster on chromosome 13 have non-synonymous substitutions in 13 sites, seven that are shared (found in more than six genes), and six that are found in only one or two genes. This variation creates 17 distinct variants of the RXLR protein encoded by this cluster (Figure 7C). Most of this variation is found in the N-terminus and only two are found in the C-terminus, which is assumed to be the functional domain that exerts its activity inside of the plant cell (Zheng *et al*., 2014). Despite extensive copy-number differences, only two out of the 23 RXLR effector orthogroups in this cluster have dN/dS values > 1 (average dN/dS = 0.57). When we consider all effecter genes in this cluster, then 14 out of the 114 effectors show positive selection (dN/dS > 1), suggesting that the expansion of effector copy-number is the leading cause of variation.

Larger sequence variation in RXLR clusters is caused by the insertion of TEs, shown as alternative paths in the pangenome graph (Figure 7A). In the first RXLR cluster, we discovered a deletion of an LTR *Gypsy* element and an RXLR gene expansion that occurred specifically in *Pe4*. The second RXLR cluster has extensive variation caused by insertion of LTR *Gypsy* elements in *Pe16*, *Pe11*, and *Pe14* that coincides with copy-number expansion of the RXLR genes. To explain the evolution of this region, we propose the following sequence of events based on the analyses of the pangenome graph. First, in the last common ancestor of isolates *Pe11*, *Pe14*, and *Pe16* two LTR *Gypsy* families were inserted around an effector gene. Second, in the last common ancestor of *Pe11* and *Pe14* the region of the two LTR Gypsies surrounding the effector gene were duplicated, thereby creating a new copy of the gene, unique for these two isolates. Third, in the ancestor *Pe16* two more LTR *Gypsy* copies were inserted upstream of the previous event (Figure 7D). The new effector copies in *Pe11* and *Pe14* are identical in sequence and share unique deletions of ten codons in centre of the gene sequence, and a unique nucleotide deletion at the 3’ end that causes an early stop. The phylogeny of the all the effector genes in the cluster shows that the subgroup of effectors, with downstream effector genes in *Pe11* and *Pe14*, is the origin of these new effector copies (S. Figure 10). These observations suggest that the activity of two LTR *Gypsy* families in this region caused the expansion of the effectors in *Pe14* and *Pe16*. Although this event explains only a part of the overall gene copy-number variation observed in this cluster, it represents a recent example of the TE insertion and gene duplication in these variable gene regions.

**Figure 7.**
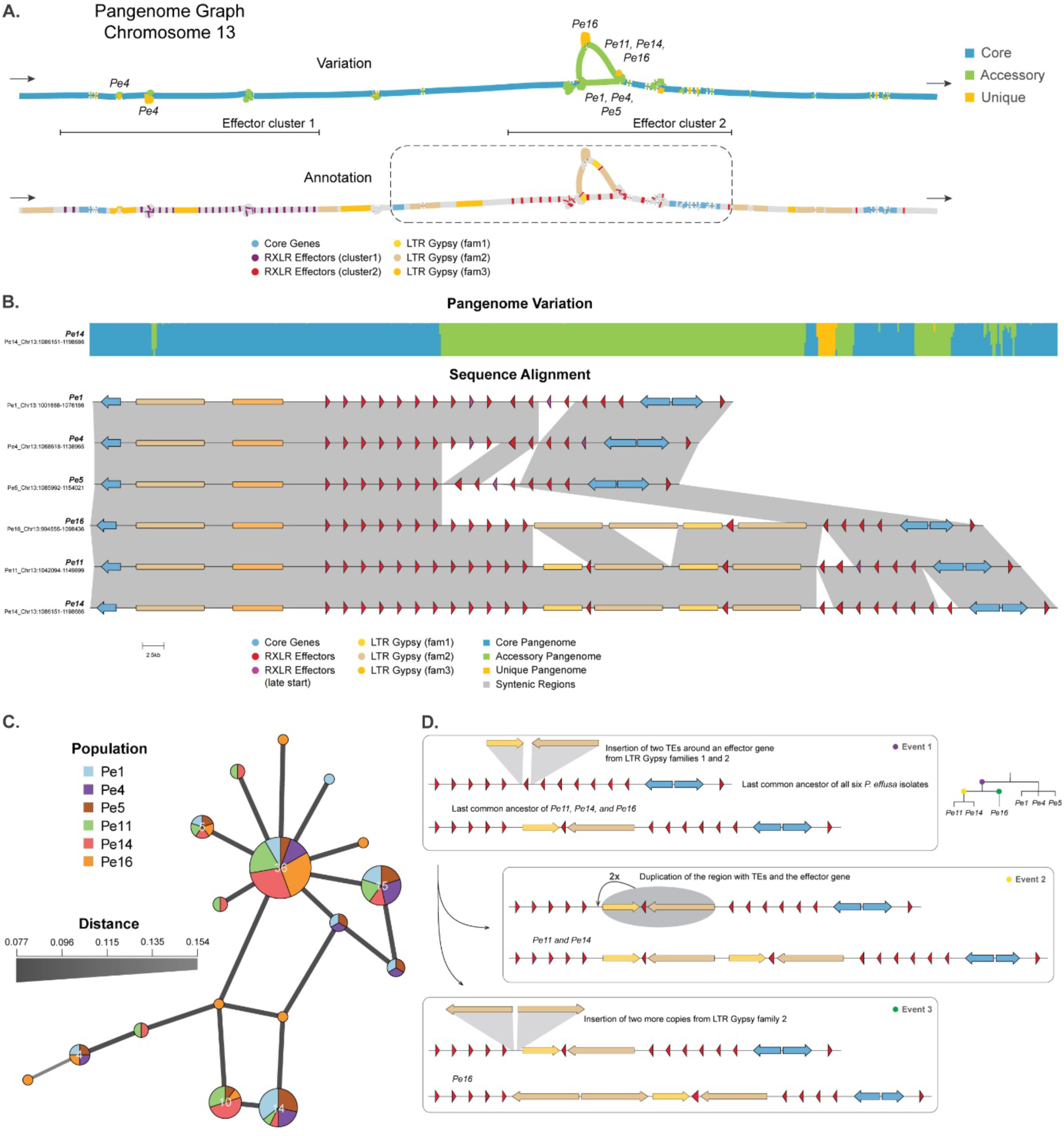
Insertions of transposable elements causes variation in effector clusters and are associated with changes in effector copy-numbers. **A.** Part of chromosome 13, 0.2 Mb in size, is visualised by two pangenome graphs, with the start and end of the graph indicated with arrows. The first graph shows the variation between *P. effusa* isolates with core (blue), accessory (green), and unique (orange) regions. For large accessory and unique regions, the isolates that have these regions are indicated. The second graph visualises the effector genes belonging to the first (purple) and second (red) effector clusters on the graph. The highlighted area is expanded in **B.** Alignment of corresponding chromosome 13 region for the six *P. effusa* isolates containing the second effector cluster. Effector genes (red), pseudogenes (purple), and core genes (blue) are indicated as arrows, and LTR Gypsy repeats (fam1: yellow, fam2: brown and, fam3: orange) are indicated as boxes. The pangenome variation for this region is visualised in a stacked bar plot (top track; core blue, accessory green, and unique orange), and syntenic regions between *P. effusa* isolates are connected with grey ribbons (bottom). **C.** Genotype network indicating the genetic variation between the RXLR effector proteins in the cluster. Each node is an effector genotype present in the second cluster on chromosome 13 and each edge is the protein sequence distance between the nodes. The size of the nodes represents the total number of effectors, and the colours of the nodes represent how each genotype is divided between the isolates. **D.** Proposed sequence of events that generated the genetic variation observed in the effector cluster across the six *P. effusa* isolates.

## Discussion

Despite extensive efforts in disease management and control, filamentous plant pathogens are rapidly evolving and continue to successfully infect their hosts (Hartmann *et al*., 2017). Pathogen adaptation is thought to be driven by few highly variable genomic regions that harbour most of the known variation in a species (Raffaele and Kamoun, 2012; Dong, Raffaele and Kamoun, 2015; Frantzeskakis, Kusch and Panstruga, 2019; Torres *et al*., 2020). Traditional comparative genomics methods, based on single reference genome sequences and short-read sequencing data, cannot fully reconstruct these regions, and thus fail to capture their variation and evolution (Everhart, Gambhir and Stam, 2021). Here we generated, to our knowledge, the first pangenome graph for an oomycete by incorporating six genetically diverse *P. effusa* isolates to improve structural annotation and to perform in-depth, genome-wide comparisons of their genes, TEs, and effectors. Our analysis revealed a highly conserved genome structure, with 80% of the observed variation being associated with TEs. Genes are typically conserved, but pathogenicity-related genes are highly variable. This variation is mainly caused by gene copy-number changes that occur inside large gene clusters, which appear to be the major mode of genome evolution in *P. effusa*. By constructing a pangenome graph, we’ve gained invaluable insights into the evolution of this important spinach pathogen. This approach can be applied to many more filamentous plant pathogens and thus presents an exciting new route to study the full extent of pathogen adaptation by precisely documenting genomic variation.

The use of pangenome graphs to compare multiple genomes has recently gained momentum in multiple organisms, including bacteria, animals, fungi, and human (Gao *et al*., 2023; Rice *et al*., 2023; Yang *et al*., 2023; Garcia *et al*., 2024). The alignment of multiple genomes of individuals or strains in a pangenome graph enables researchers to construct a common structural annotation, thus consistently discovering genes for each genome and reducing potential variation that could have been merely caused by different technical interpretations in the separate structural annotation of the individual genomes (Fiddes *et al*., 2018). For example, we identified in total 1,842 genes that would have been missed in the individual annotations (S. Note 2). We also demonstrate that the pangenome graph can be used to directly determine the synteny of genes and to create synteny-anchored single-copy orthogroups. Based these pangenome based methods of structural annotation, the full extent of gene variation between the genomes can be discovered, including gene copy-number variations.

Large-scale chromosomal rearrangements between isolates are commonly observed in filamentous fungi such as in *Verticillium dahliae* and *Magnaporthe oryzae*, and these have been proposed to be important to generate variation in the absence of sexual recombination (De Jonge *et al*., 2012; Seidl and Thomma, 2014; Faino *et al*., 2016; Menardo *et al*., 2017; Van Dam *et al*., 2017; Hoogendoorn *et al*., 2018; Langner *et al*., 2021; Westerhoven *et al*., 2024). In contrast, chromosome structure is highly conserved in *P. effusa*, even though we selected phylogenetically distant isolates for our analyses. The conserved chromosome structure is largely shared with other species within the Peronosporaceae such as *Bremia lactucae* and *Peronosclerospora sorghi*, with the exception of few chromosomal fusions (Fletcher and Michelmore, 2023). Our analysis indicates that the chromosomes 1 and 10 might also be the result of a fusion of two smaller chromosomes, showing that chromosome fusions indeed do occur in Peronosporaceae. Nevertheless, they show a remarkable chromosomal stability that could result from sexual reproduction that is an essential part of the yearly life cycle of most oomycetes (Feng *et al*., 2020; Skiadas *et al*., 2022).

In contrast to the overall conserved chromosome structure in Peronosporaceae (S. Figure 5A), we discovered the presence of an accessory chromosome in 19 out of 32 *P. effusa* isolates that is does not correspond with any of the chromosomes in *B. lactucae* or *P. sorghi* (Fletcher *et al*., 2021, 2023). The accessory chromosome is highly repetitive, and the activity of LINEs and LTRs appears to underlie the variation observed between *P. effusa* isolates (S. Figure 7; Panel Pe5_Chr18). Despite their activity, none of the TE families found on chromosome 18 can be found on any other chromosome, except for two LTR families which experienced recent expansion following the differentiation of the *P. effusa* isolates. Consequently, it seems more likely that this chromosome was recently acquired from a close relative, possibly from *P. farinosa* f. sp. *betae*, rather than deleted from multiple isolates. Accessory chromosomes in asexually reproducing filamentous fungi, like *Fusarium oxysporum*, are often associated with pathogenicity (Hartmann *et al*., 2017; Van Dam *et al*., 2017; Westerhoven *et al*., 2024), but since we were not able to identify protein-coding genes on chromosome 18, its physiological significance remains uncertain. Additionally, we observed degradation of this chromosome in many isolates that likely emerged from sexual reproduction between *P. effusa* isolates (Figure 4D) (Skiadas *et al*., 2022), suggesting that sexual reproduction could be contributing to its degradation.

By creating chromosome-level genome assemblies based on long-read data, we were able to discover the TE content and its variation in *P. effusa*. Incomplete genome assemblies typically miss regions of recent TE activity (Thomma *et al*., 2016), which for example explains the discrepancy in genome sizes compared with previous *P. effusa* reference genomes (28 Mb larger) (Fletcher *et al*., 2018; Klein *et al*., 2020). Our pangenomic comparison reveals that the TE expansion in *P. effusa* is, in many cases, more recent than the differentiation of the here analysed isolates. Most notably, SINE-like sequences have recently expanded and contribute to the extensive variation between the *P. effusa* isolates. TE expansion in other oomycete genomes has been thought to play a similar role in their genome expansion and variation (Haas *et al*., 2009; Raffaele and Kamoun, 2012; Faino *et al*., 2016). Notably, while other TE families have been expanding and recent copies can be found on multiple chromosomes, each SINE-like family in *P. effusa* is specific to a single chromosomal location, and expansion creates large repetitive regions of almost identical sequences. Additionally, it is worth noting that these elements are commonly annotated as tRNA. For example, the recent annotation of more than 7,000 tRNA genes in *Phytophthora infestans* suggests that these SINE-like elements are active and play an important role in oomycete genome evolution (Matson *et al*., 2022).

The pangenome graph uncovered extensive and recent changes in gene copy-number, especially for RXLR and CRN effectors as well as for other pathogenicity-related genes. While the molecular mechanism has yet to be discovered, recent TE activity is often observed close to genes that display gene copy-number variation. This phenomenon has also been described in *Phytophthora*, where effector gene copy-number variation was shown for RXLR genes *Avr1a* and *Avr3a in Phytophthora ramorum* (Mathu Malar *et al*., 2019) and RXLR and CRN genes in *Phytophthora sojae* (Qutob *et al*., 2009), and these copy-number changes are thought to impact pathogen fitness (Qutob *et al*., 2009; Mathu Malar *et al*., 2019). It is also well established that NLP genes have recently expanded in oomycetes by gene duplications and that these genes are organised in clusters (Seidl and Van Den Ackerveken, 2019). In *P. effusa* 15/18 NLPs are organised in two clusters in which all the variation between NLPs can be observed. Interestingly, when comparing the orthologous genes, we observed that clustered effectors more often evolve under positive selection than unclustered effectors, suggesting that locally copied effectors rapidly generate novel genotypes. Thus, dynamic gene clusters, embedded in otherwise highly conserved chromosomes, evolve by extensive copy-number changes and serve as cradles for genomic variation in oomycetes. Given the frequency and the emergence of novel *P. effusa* races, copy-number variation of pathogenicity genes and diversification of novel gene copies, likely plays an important role in the adaptation of *P. effusa*. Adding more and more closely related *P. effusa* isolates to the pangenome will therefore be essential to associate effector variation with virulence and the capacity to break deployed spinach resistance traits.

Pangenome graphs will most likely transform the field of comparative genomics, resulting in more accurate and in-depth analysis of the variation of a large collection of diverse genomes (Badet and Croll, 2020; Everhart, Gambhir and Stam, 2021). Especially chromosome-level genome assemblies will be essential to achieve the necessary resolution to uncover and study highly variable regions that are enriched for pathogenicity-related genes (Fletcher and Michelmore, 2023; Westerhoven *et al*., 2024). Beyond adding genomes to the graphs, a step further would be to create a pangenome graph that incorporates fully phased diploid or polyploid genome assemblies, thus uncovering possible variation and recombination between haplotypes (Henningsen *et al*., 2024). For example, oomycetes are mostly diploids, but there are various reports of whole-genome duplication in *Phytophthora betacei*, and aneuploidy in *Phytophthora capsici* and in *Phytophthora cinnamomi* (Seidl *et al*., 2012; Kasuga *et al*., 2016; Hu *et al*., 2020; Ayala-Usma *et al*., 2021; Engelbrecht *et al*., 2021). We therefore anticipate that pangenomic approaches will be instrumental to uncover the full extent of genomic variation in filamentous plant pathogens (Sirén *et al*., 2021; Garcia *et al*., 2024), which, in turn, will unveil new theories about their emergence and evolution, impacting our ability to predict and manage plant diseases.

## Materials and methods

### *Peronospora effusa* infection on soil-grown spinach and spore isolation

Spinach plants were sown in potting soil (Primasta, the Netherlands) and kept under long-day conditions (16-hour light, 21 °C). Two to three weeks after germination, the spinach plants were inoculated with *P. effusa* by spraying them with *P. effusa* spores suspended in water using a spray gun. Following inoculation, we placed the plants under 9-hour light and 16 °C, and the lids of the plastic trays were sprayed with water and covered to keep the plants humid. After 24 hours, the vents on the lids were opened. The lids of the boxes were again sprayed with water and the vents were closed 7-10 days after inoculation, creating a humid environment that promotes the sporulation of *P. effusa*.

To harvest *P. effusa* spores for Oxford Nanopore sequencing, we collected leaves with sporulating *P. effusa* from spinach plants and placed them in a glass bottle with tap water. The spores were brought into suspension by shaking the bottle vigorously. Soil and other large contaminants were removed by filtering the spore suspension over a 50-μm nylon mesh filter (Merck Millipore, USA). To remove small biological contaminants, the remaining filtrate was filtered 11-μm nylon mesh filter (Merck Millipore, USA) using the Merck™ All-Glass Filter Holder (47 mm) and a vacuum pump, resulting in the spores which remaining on top of the filter and contaminants washing through the filter. The spores were washed several times, scraped off the filter, and kept at -80 °C.

### High-molecular weight DNA extraction protocol

To isolate high-molecular weight (HMW) DNA, the collected *P. effusa* spores were ground to a fine powder in liquid nitrogen together with 0.17-0.18mm glass beads. The ground spores were washed in cold Sorbitol solution (100 mM Tris-HCl pH 8.0; 5 mM EDTA pH 8.0; 0.35 M Sorbitol, 1% PVP-40, 1% β-mercaptoethanol, pH 8.0). The tissue was lysed by incubation in extraction buffer (1.25 M NaCl, 200 mM Tris.HCl, pH 8.5, 25 mM EDTA, pH 8.0, 3% CTAB, 2% PVP-40, 1% β-mercaptoethanol) and incubated with proteinase K and RNase A for 60 minutes at 65 °C, and throughout the incubation mixed by gentle inversion. The samples were centrifuged to pellet and remove the debris. HMW DNA was further purified by phenol/chloroform/IAA and chloroform/IAA extraction, another RNase treatment, extraction with phenol/chloroform/IAA and chloroform/IAA and isopropanol precipitation. DNA concentration and integrity were determined using Nanodrop, Qubit, and Tapestation.

### Genome sequencing using Oxford Nanopore

We obtained long-read sequencing data for six *P. effusa* isolates (*Pe1*, *Pe4*, *Pe5*, *Pe11*, *Pe14* and *Pe16*) with Oxford Nanopore sequencing technology (Oxford Nanopore, UK) at the USEQ sequencing facility (the Netherlands). The ligation-based sequencing kit from Oxford Nanopore was used for library preparation (ONT - SQK-LSK109; Oxford Nanopore, UK) following the manufacturer’s protocol. We used a Nanopore MinION flowcell (R10) for real-time sequencing and base-calling of the raw sequencing data was performed using Guppy (version 4.4.2; default settings). The raw long-read sequencing data were checked for contamination using Kraken2 (version 2.0.9; default settings) (Wood, Lu and Langmead, 2019).

### Genome sequence using Hi-C

We obtained Hi-C sequencing data for six *P. effusa* isolates (*Pe1*, *Pe4*, *Pe11*, *Pe14*, and *Pe16*). Spores were crosslinked in formaldehyde 1% and incubated at room temperature for 15 minutes with periodic mixing. Glycine was added to a final concentration of 125 mM, followed by incubation for another 15 minutes. The spores were pelleted by centrifugation and washed with Milli-Q water. Glass beads were added, and the samples were vortexed. Crosslinked samples were pelleted by centrifugation, kept at minus 80 and send to Phase genomics (USA) for sequencing.

### Transcriptome sequencing using RNAseq

RNA was extracted from spores or infected spinach leaves with a Kingfisher System using the MaxMag Plant RNA isolation Kit. The sequencing was done at USEQ (the Netherlands), using the Truseq RNA stranded polyA library prep and the samples were sequenced on the NextSeq500 platform with 2 x 75 bp mid output (210 M clusters).

### Genome assembly

We used the long-read Oxford Nanopore sequencing data to produce chromosome-level genome assemblies for six *P. effusa* isolates. The reads were corrected, trimmed, and assembled using Canu (version 2.3) (Koren *et al*., 2017) with the following command:

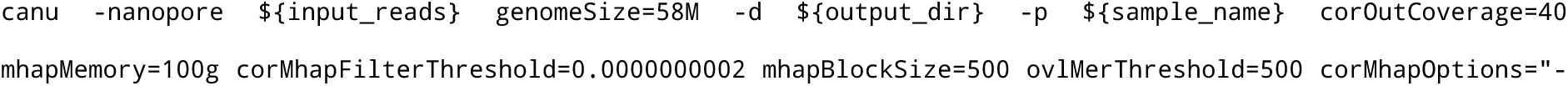
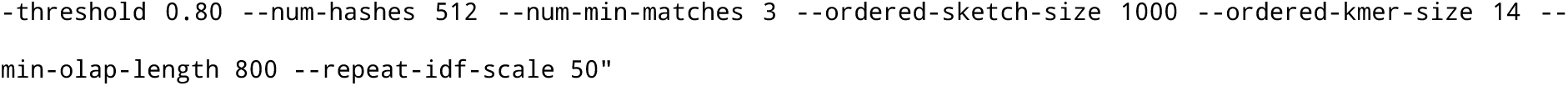

To remove contigs that come from possible contaminants, uncollapsed haplotypes, and other assembly artefacts, we created a pipeline to filter and curate draft genome assemblies. First, uncollapsed contigs were removed with Purge Haplotigs (version 1.1.1; -a 90) (Roach, Schmidt and Borneman, 2018). The taxonomy of the remaining contigs was determined with CAT (version 5.2; default settings) (Von Meijenfeldt *et al*., 2019) and was visualised together with the Nanopore read coverage and QC content with Blobtools (version 2.3.3; default settings). The contigs classified as contamination were removed, while contigs classified as oomycete, Peronosporaceae, *Peronospora*, and the unclassified contigs were retained. The mitochondrial contigs were removed based on their characteristically different GC content (22% GC of the mitochondrion vs 44-52% of the nuclear genome for *P. effusa*). Finally, genome assembly metrics were measured with QUAST (version 5.0.2) (Mikheenko *et al*., 2018).

### Scaffolding assemblies with Hi-C data and closing gaps

The curated assemblies were further scaffolded to full chromosomes by using Hi-C short-read data that were aligned to the reference assemblies with Juicer (version 1.6; default settings) (Durand, Shamim, *et al*., 2016). Based on this alignment, the pairwise Hi-C contacts along the contigs were visualised on a heatmap generated with 3D-dna (version 180922, -r 0 -e) (Dudchenko *et al*., 2017). Based on the heatmap, the contigs were manually scaffolded to 17 or 18 chromosomes using Juicebox (Normalisation: balanced, Resolution: 50Kb) (Durand, Robinson, *et al*., 2016).

Gaps of the scaffolded chromosomes were closed with FinisherSC (version 2.1; default settings) (Lam *et al*., 2015). The scaffolded assemblies were corrected for single nucleotide polymorphisms using Illumina short-reads with four rounds of Pilon (version 1.23; --diploid, --fixbases) (Walker *et al*., 2014). The Illumina short-read data was published previously by Skiadas et al. (Skiadas *et al*., 2022).

### Transposable element annotation

To create a combined transposable element (TE) library for all *P. effusa* isolates, we used EarlGrey (version 2.0) (Baril, Galbraith and Hayward, 2024), with dependencies RepeatMasker (version 4.1.2) (Smit, Hubley and Green, 2013), RepeatModeler (version 2.0.2) (Flynn *et al*., 2020), and DFAM (version 3.6) (Storer *et al*., 2021). First, the EarlGrey pipeline was run on the genome of *Pe1* with option ‘-r 2759’, specifying the subset of eukaryotic sequences in the DFAM database as reference TE library. The TE library produced was then expanded by using it as an input for the EarlGrey pipeline, rather than the DFAM library, and recursively running it with additional *P. effusa* isolates (S. Figure 12). TE consensus sequences with hits of 0.001 e-value or lower, at least 30 matching positions and a coverage and identity of at least 80% were discarded. The final combined TE library was then filtered for TE families that overlapped with gene annotations that have also RNAseq coverage on the *P. effusa* assembly, thus removing 21 consensus sequences from the library. This combined and filtered TE library was then used to annotate and soft-mask the genomes using RepeatMasker (version 4.1.2 -e rmblast -xsmall -s -nolow).

### Genome annotation

The soft-masked genomes and the RNAseq short-read data were used for structural gene prediction and functional annotation with the funannotate pipeline (version 1.8.7) (Palmer, 2017) with the following commands:

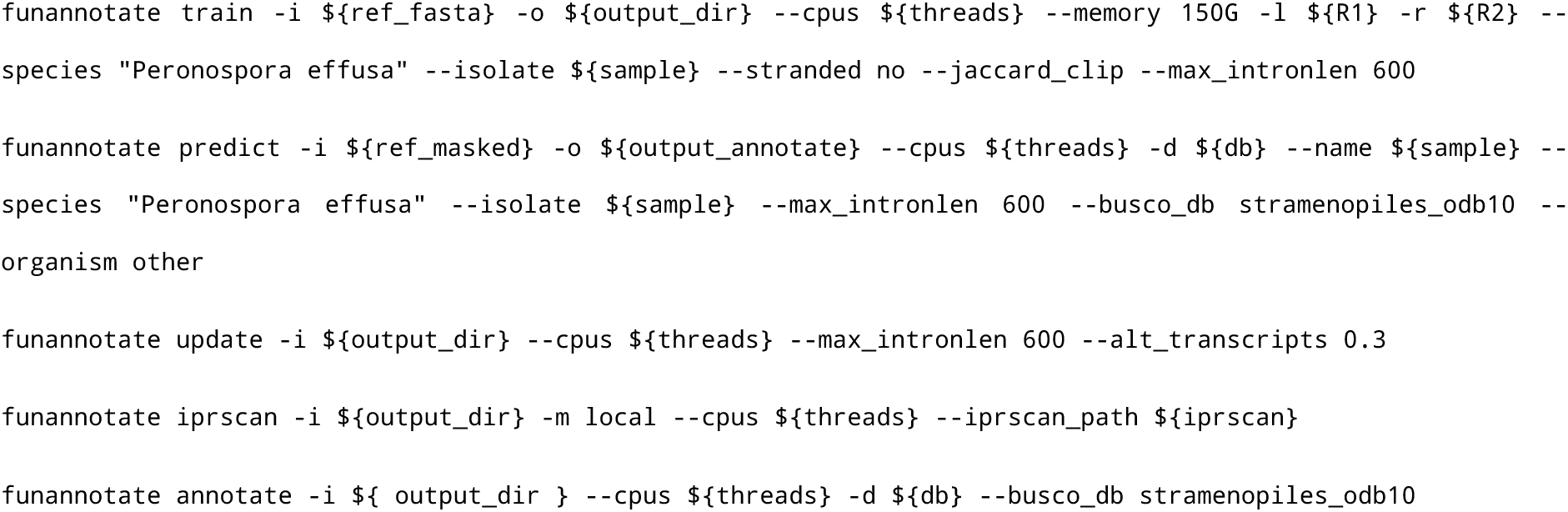

The assembled and structurally annotated genomes were visualised with Circos (v. v 0.69-8) (Krzywinski and Ave, 2015) (Figure 1A).

### Secretome and effector prediction

In addition to genes annotated by funannotate, we also extracted all open reading frames longer than 70 amino acids encoded in the repeat-masked genome using ORFfinder (v.0.4.3, -ml 210 -s 0). We filtered ORFs when they overlapped with the annotated genes using bedtools intersect (version 2.30.0, default settings) (Quinlan and Hall, 2010).

For the prediction of the secretome of *P. effusa* races, we used the Predector pipeline (v. 1.2.6, default settings) (Jones *et al.,* 2021). We extracted the list of all sequences predicted to contain a signal peptide by at least one of the tools included in the pipeline (SignalP3.0, SignalP4.0, SignalP5.0, SignalP6.0, Deepsig, and Phobius) (Dyrløv *et al*, 2004; Petersen *et al.,* 2011; Almagro *et al.,* 2019; Teufel *et al.,* 2021; Kall *et al.,* 2007; Savojardo *et al.,* 2017). From those, the ones containing multiple transmembrane domains according to Phobious or TMHMM (Krog *et al.,* 2001) were excluded from the secretome.

The secreted proteins were then screened to detect the presence of the conserved motifs described in RXLR and Crinkler oomycete effectors (Saraiva *et al*., 2022). To perform regular expression searches, the EffectR package for R (Tabima and Grünwald, 2019) was used to detect the following patterns in the first 100 amino acids after the signal peptide: i) a divergent version of the RXLR-EER complete motif ([RQGH]XL[RQK]-[ED][RK] (Haas *et al*., 2009), and ii) canonical and divergent versions of the RXLR motif alone (RXLR or [RQGH]XL[RQK]). For the Crinklers, we used regular expressions to detect the occurrence of canonical LFLAK, a degenerated version with maximum two allowed changes in the motif (established as L[FYRL][LKF][ATVRK][KRN] by Zhao *et al*., 2020), and the HVLV motif.

We also performed sequence profile searches using HMMER v3.3 (Mistry *et al*., 2013). To create the sequence profile for the WY domain (Boutemy *et al*., 2011), the sequences described were aligned using MAFFT v7.453 (ginsi) (Katoh *et al*., 2002), and an HMM profile was built from the alignment using hmmbuild. This profile was used to screen the secreted proteins using hhmsearch (--incE 10). A similar approach was followed to create profiles to search for the RXLR(-EER) motifs using the 253 reviewed RXLR effectors deposited at the UniProt database (extracted with the search string “family:“rxlr effector family” AND reviewed:yes” in January 2022) (Bateman *et al.,* 2020). Three different profiles were built from the proteins annotated to contain a RXLR motif, an EER motif, or both RXLR-EER. To search for Crinklers, we used an HMM profiles for the LFLAK and the DWL domains described by Armitage *et al*. (2018).

The list of RXLR candidates includes all secreted proteins displaying a canonical or divergent RXLR-EER motif, the canonical RXLR, the WY domain, or a divergent RXLR, in case the protein was also found by least one of the Uniprot HMM profiles. For Crinklers, all proteins identified with the LFLAK/DWL regular expressions or HMM profiles were included in the list of candidate effectors.

### Pangenome graphs and common annotation

A pangenome graph was built per chromosome by aligning the genome assemblies using Minigraph-Cactus (version 2.6.4, --filter 0 --vcf clip full --gfa clip full) (Hickey *et al*., 2023). Pangenome graphs from minigraph were visualized with bandage (version 0.8.1) (Wick *et al*., 2015).

The hal output of the pangenome graph, the gene annotation, and the RNAseq coverage for each *P. effusa* isolate were used to collectively reannotate genes using evidence from all isolates with the Comparative-Annotation-Toolkit (version 2.2.1, --augustus --augustus-cgp --assembly-hub --filter-overlapping-genes) (Fiddes *et al*., 2018). Genes that were not characterized as protein-coding or had no alternative source transcripts were removed from the annotation (gene_biotype=protein_coding).

### Gene evolution

To calculate the dN/dS ratio, we used the python package dnds (version 2.1). The calculation was made pairwise for each ortho-group, excluding those that would result in a division by zero or with dN/dS score higher than five to avoid saturation of substitutions (De La Torre *et al*., 2017).

### Splits tree

We performed variant calling, using the genome assembly of *Pe1* as a reference, and the short-read data of 32 *P. effusa* isolates, and generated a joint VCF file with both variant and invariant sites with GATK (version 4.4.0.0, GenotypeGVCFs -all-sites). The single nucleotide variants of were transformed into a distance matrix with PGDSpider (version 2.1.1.5) (Lischer and Excoffier, 2012), which was then was used to construct a decomposition network using the Neighbor-Net algorithm with SplitsTree (version 4.17.0) (Huson and Bryant, 2006). We calculated the branch confidence of the network using 1,000 bootstrap replicates.

### Effector genotype network

We collected from all six *P. effusa* isolates the effector protein sequences from the effector cluster in chromosome 13 and we performed variant calling using GATK (version 4.4.0.0). The variant sites were used to create a genotype network using to R package clusterPoppr (version 2.9.4, default settings) (Kamvar, Tabima and Gr̈unwald, 2014).

### Visualisations

Using ModDotPlot (version 0.7.2) (Sweeten, Schatz and Phillippy, 2024), we visualised the repeat in chromosome 18 (-id 75 -k 11 -r 3000) and the repetitive region in chromosome 6 (-id 85 -k 11 -r 500). Clinker (version 0.0.28) (Gilchrist and Chooi, 2021) was used to visualise the effector cluster on chromosome 13, by aligning the gene protein sequences of six *P. effusa* isolates in that region. All other visualisations were created in python using matplotlib and seaborn (Waskom, 2021).

## Supporting information

STable1

STable2

STable3

## Author Contributions

PS and MFS conceived and designed the data analysis. JE and MNM generated the data. PS, SRV, and JD performed the data analysis. PS visualised the results and wrote the paper. MFS, RDJ, and GVDA acquired funding for this project, supervised, and reviewed the paper.

## Acknowledgments

We acknowledge the Utrecht Sequencing Facility (USEQ) for providing sequencing service and data. USEQ is subsidized by the University Medical Center Utrecht and The Netherlands X-omics Initiative (NWO project 184.034.019).

## Data availability

For this study, we sequenced isolates of the 19 denominated races of *P. effusa*. Cultures of these can be requested for research from Naktuinbouw, the Netherlands (Correll *et al*., 2015) (S. Table 3). The here as well as previously generated raw sequence data and genome assemblies generated in this study are available at NCBI under BioProject PRJNA772192. The code written for this study is available in github https://github.com/Umbel89/pangenome_analysis.

## Supplementary Notes

Supplementary Note 1: Chromosome-level genome assemblies for six *Peronospora effusa* isolates

To obtain high-quality, chromosome-level genome assemblies, we produced on average, 12 Gb of Nanopore reads per *Peronospora effusa* isolate, with an N50 of 28.5 kb, and 8 Gb of Hi-C reads (S. Table 1). The Nanopore long-read data were assembled with Canu (Koren *et al*., 2017), and assembled contigs were subsequently filtered to remove the mitochondrial genome and bacterial contamination (S. Figure 2B), which is a known challenge for biotrophic plant pathogens (Strong *et al*., 2014; Klein *et al*., 2020), yielding 29-100 contigs per isolate. These contigs were further scaffolded into chromosomes using the chromatin contact information from Hi-C data (S. Figure 4), and remaining gaps were subsequently closed by manually joining contig overlaps that were supported by long reads. To correct for systematic sequencing errors of the Nanopore data, the chromosome-level genome assembly was polished with Illumina short-read data.

The quality and completeness of the assembly was evaluation in a reference free method, using short-read K-mer distribution and observed that our haploid assembly has the expected coverage in the heterozygous (1x), homozygous (2x), and highly repetitive regions (>3x) (S. Figure 11). We further evaluated the quality of the genome assemblies based on the coverage of Nanopore and Illumina reads along the assembled chromosomes, which support the chromosome structure with average coverage per isolate ranging from 53 - 271 reads (S. Figure 3); the number and constitution of the assembled chromosomes is also strongly supported by the Hi-C data (S. Figure 4). We did, however, also observed lower sequencing coverage for few long repetitive regions (longer than 100 kb), but the contiguity of these regions was supported by Nanopore reads that were longer than the respective region (S. Figure 3). The ribosomal RNA cluster at the beginning of chromosome 15 has five times higher Nanopore read coverage compared with the remainder of the genome, suggesting that this highly repetitive region is much longer than represented in the assembly.

Supplementary Note 2: Pangenome graph of six *Peronospora effusa* isolates enables comprehensive study of genome variation

To annotate transposable elements (TEs), we applied a method for common TE discovery, where the library from the *de novo* TE discovery of *Pe1* was used to mask the genome of *Pe4*, before performing *de novo* TE discovery on *Pe4* and appending the common library with the new sequences discovered on *Pe4* (S. Figure 12). This method was recursively applied for *Pe5*, *Pe11*, *Pe14*, and *Pe16* resulting in a common TE library for all assembled isolates. This common TE library was subsequently used to annotate each genome assembly, which uncovered that between 50.58 and 52.4% of each genome is composed of repetitive sequences such as transposons, with long-terminal repeat (LTR) transposons being the most abundant superfamily covering around 40% of the genome (Figure 1C) (Materials and Methods – Transposable element annotation). We used repeat annotated genomes as a basis to structurally annotate genes with funannotate (Palmer, 2017) by incorporating RNAseq data from *Pe1*, *Pe5*, *Pe11*, *Pe14*, *and Pe16* together with *ab initio* and homology-based gene annotation, which resulted in 9,869 to 10,008 protein-coding genes and in 6,092 to 7,362 tRNA genes.

To be able to consistently annotate genes for multiple *P. effusa* isolates and to overcome potential errors introduced by the separate gene annotation, we exploited the information in the pangenome graph to reannotate genome assemblies. To this end, we used the annotations of each isolate, the available RNAseq data, and the alignment of each genome to the pangenome graph to perform a common annotation using the Common-Annotation-Toolkit (Figure 2B, steps 1-3) (Materials and Methods – Pangenome graphs and common annotation) (Fiddes *et al*., 2018; Hickey *et al*., 2023).

To compare the annotated protein-coding genes for each *P. effusa* isolate, we assigned all genes into groups based on their position on the pangenome graph, thus creating single-copy gene orthogroups based on synteny along the graph rather than the traditional approaches that group genes based on sequence similarity alone (Figure 2B, step 4). This method results in 12,379 orthogroups, of which 9,031 (73%) are conserved in the pangenome, while for each isolate 86.8 to 90.9% of the predicted protein-coding genes are conserved. Most unique genes are found in *Pe5* (4.3%, 450) and the lowest number in *Pe1* (1.1%, 110).

## Supplementary Figures

**Supplementary Figure 1.**
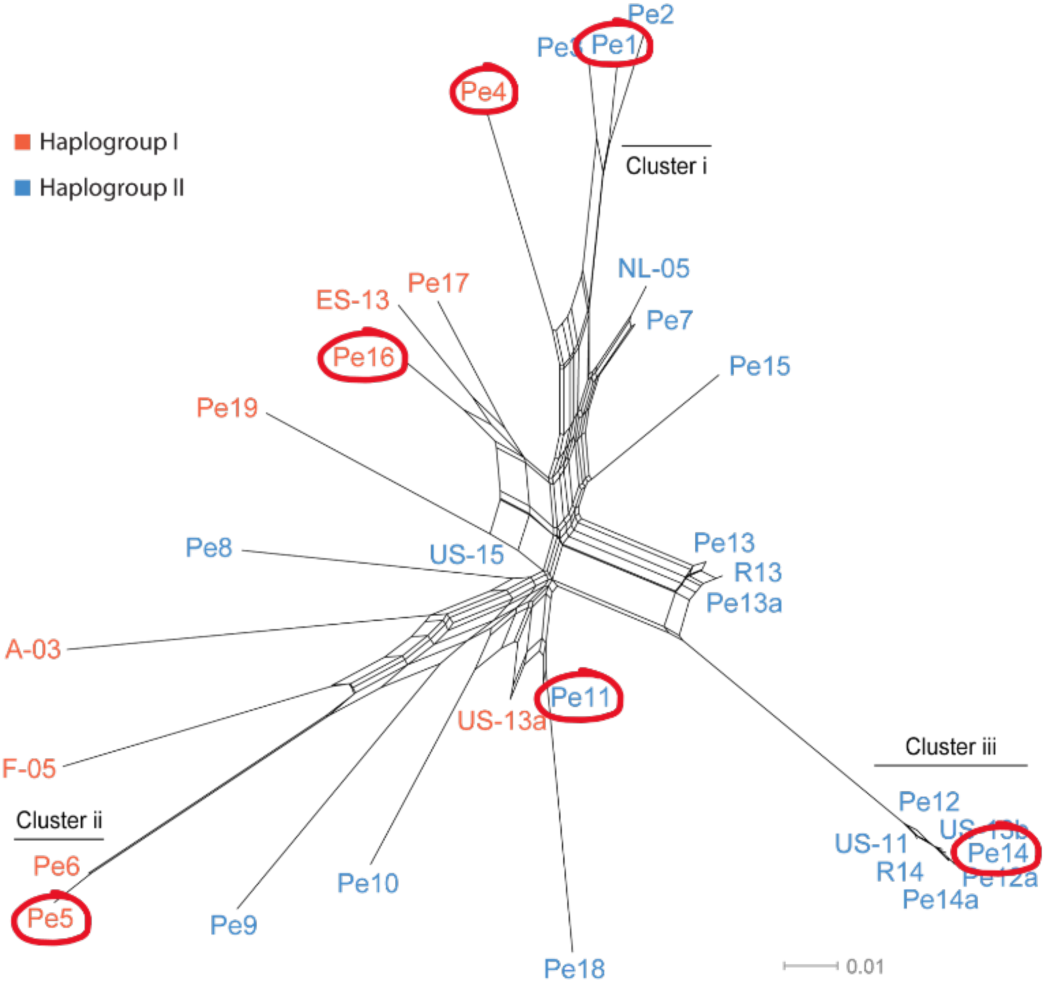
Six isolates were selected to capture the genomic variation of *Peronospora effusa*. Neighbor-net phylogenetic network of *P. effusa* isolates that cover all known isolates based on a distance matrix of genome-wide nucleotide differences, containing 260,616 biallelic sites. The branch lengths are proportional to the calculated number of substitutions per site. The parallel edges connecting different isolates indicate conflicting phylogenetic signals. The six selected isolates are from six distinct races and are indicated with red circles.

**Supplementary Figure 2.**
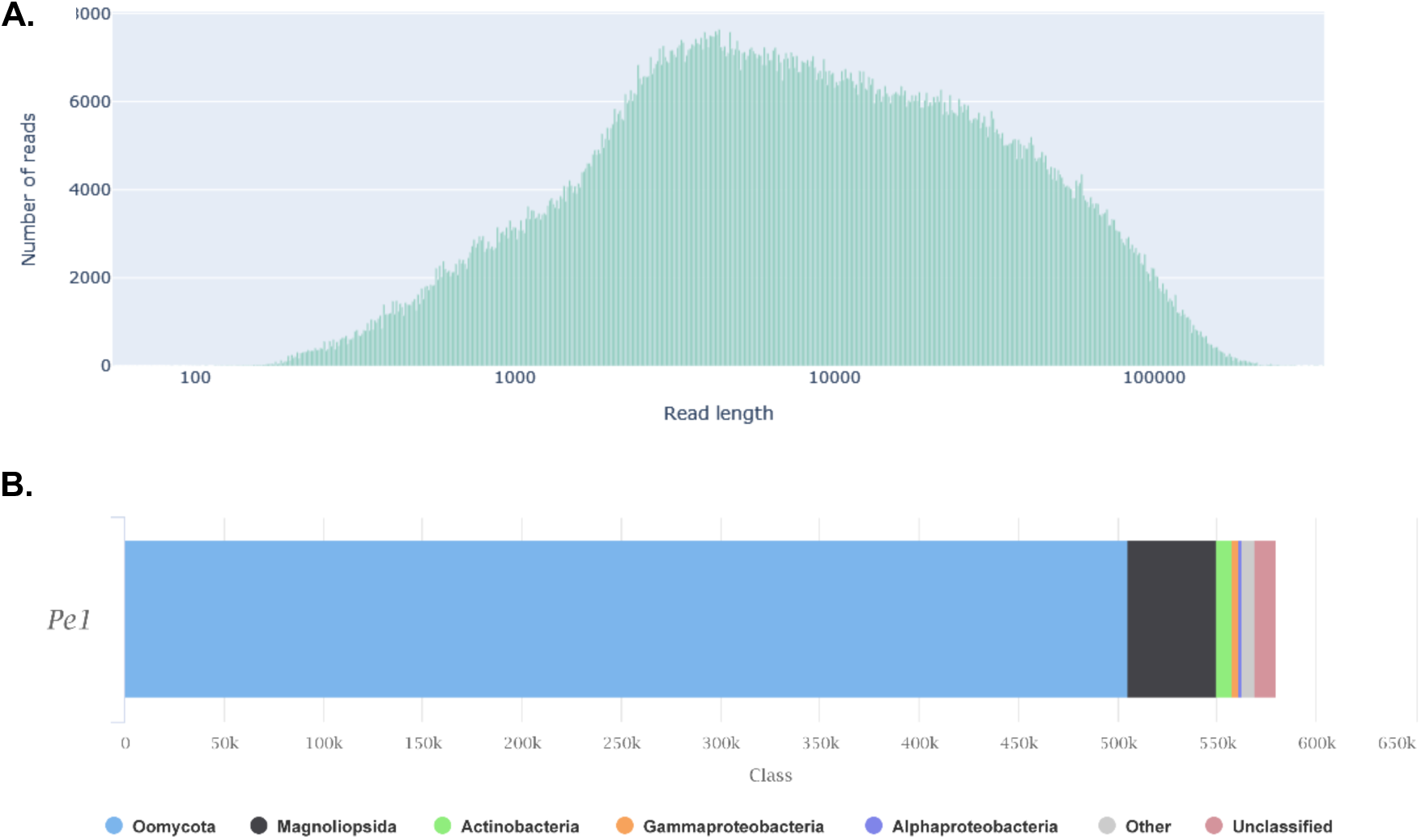
Nanopore sequencing for *Pe1* isolate resulted in high quality long-reads. **A.** Weighted histogram of read lengths after log transformation. **B.** Phylogenetic classification of reads based on Kraken2 (Wood et al., 2019) and plotted with MultiQC (Ewels et al., 2016). All the reads classified as oomycote belong to *P. effusa*.

**Supplementary Figure 3.**
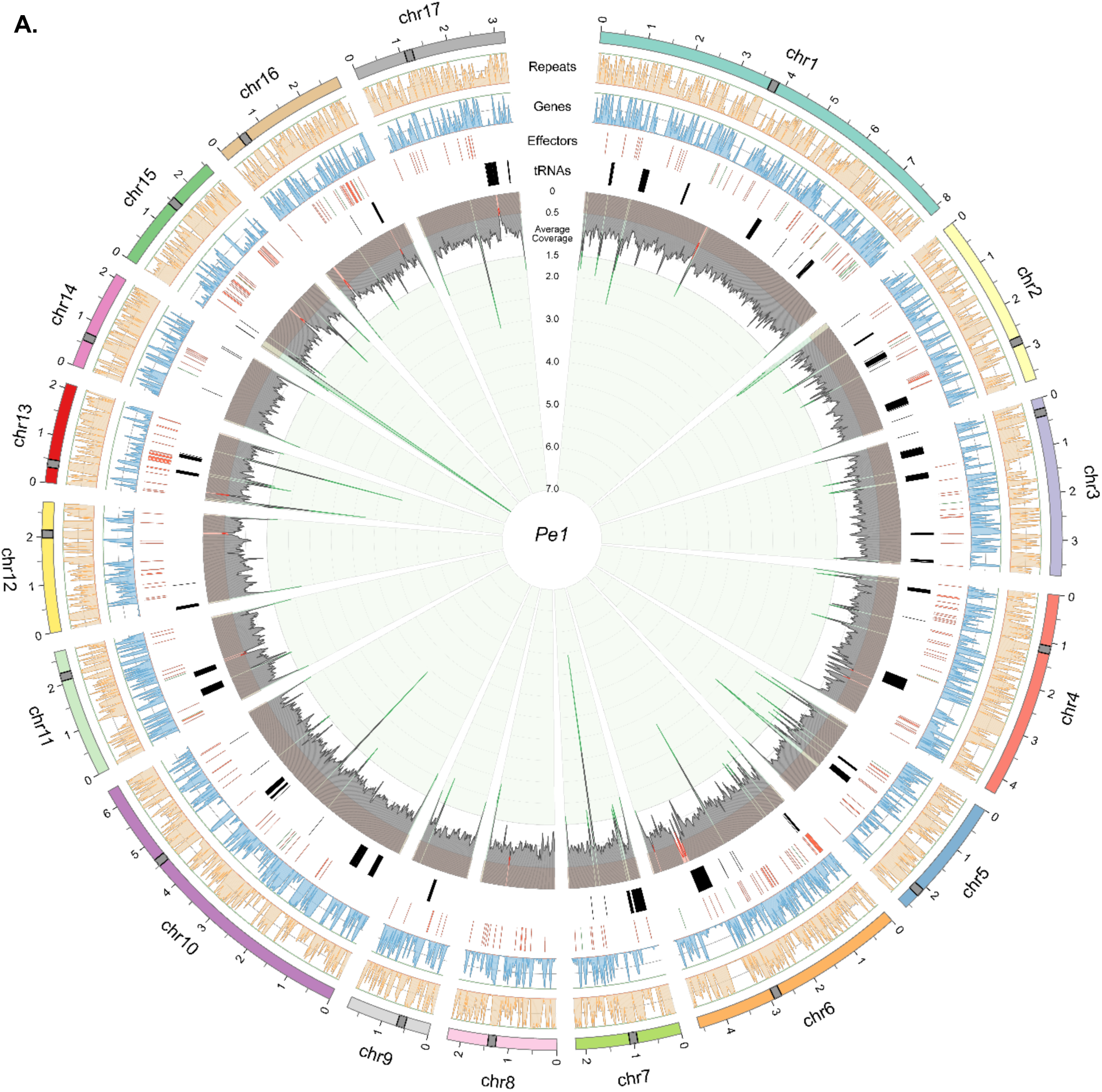
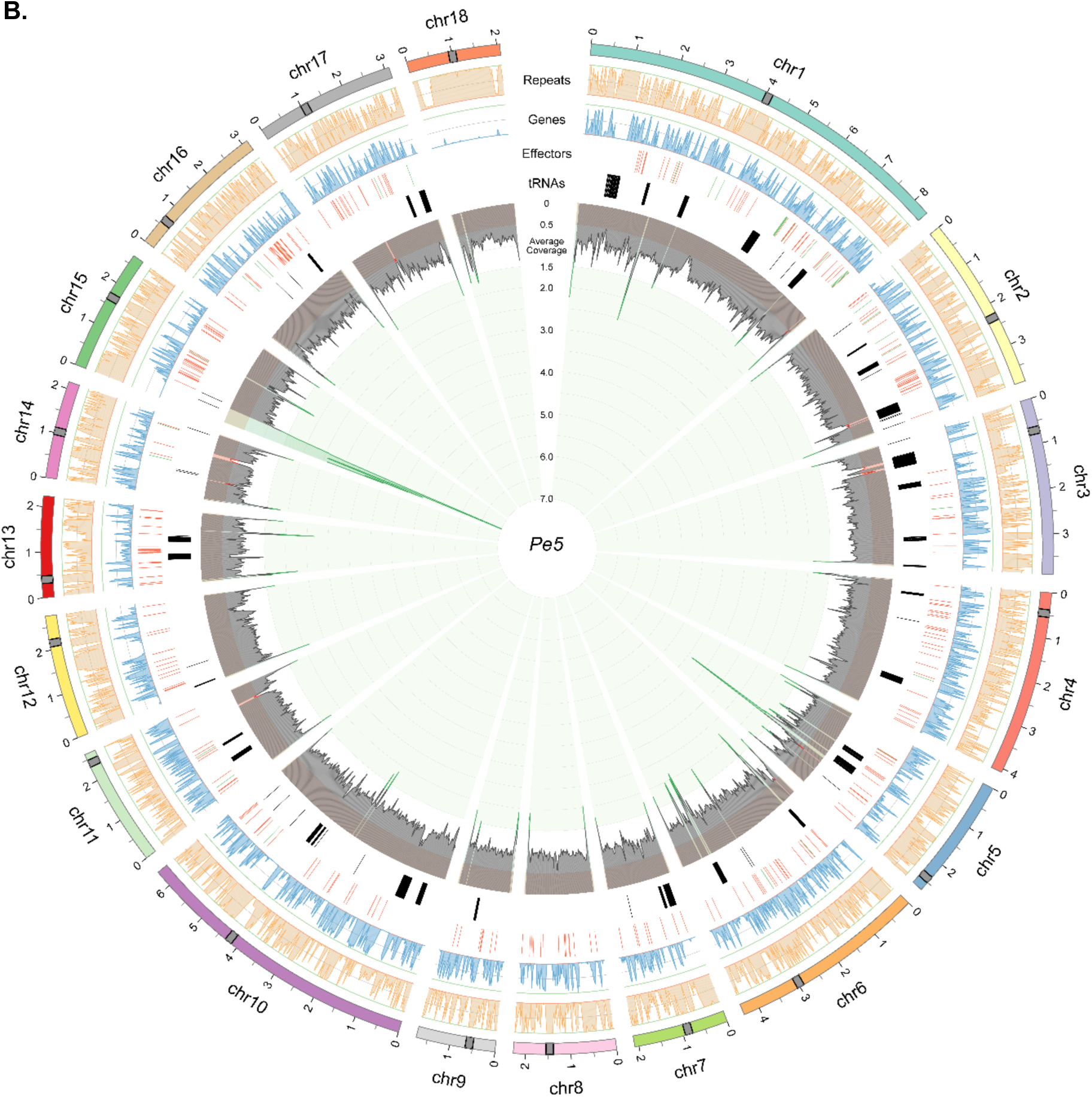
The genome assemblies of *Peronospora effusa* race 1 (*Pe1*) and race 5 (*Pe5*) are visualised in a circular plot. Individual tracks, starting from the outside to the inside: i) 17/18 chromosomes are shown in different colours, a grey rectangle point the location of the centromeres. ii) Line graph shows the coverage of repeat (orange) and gene (blue) content summarized in non-overlapping 20 kb windows. iii) Lines indicate the position of RXLR (red), CRN (green), and tRNA (black) genes. iv) Inverted line graph shows the nanopore average coverage of nanopore reads to the genome assembly. Coverage of 1 indicates a diploid coverage, 0.5 haploid coverage, and coverage higher than one indicates an underrepresentation of a repetitive region.

**Supplementary Figure 4.**
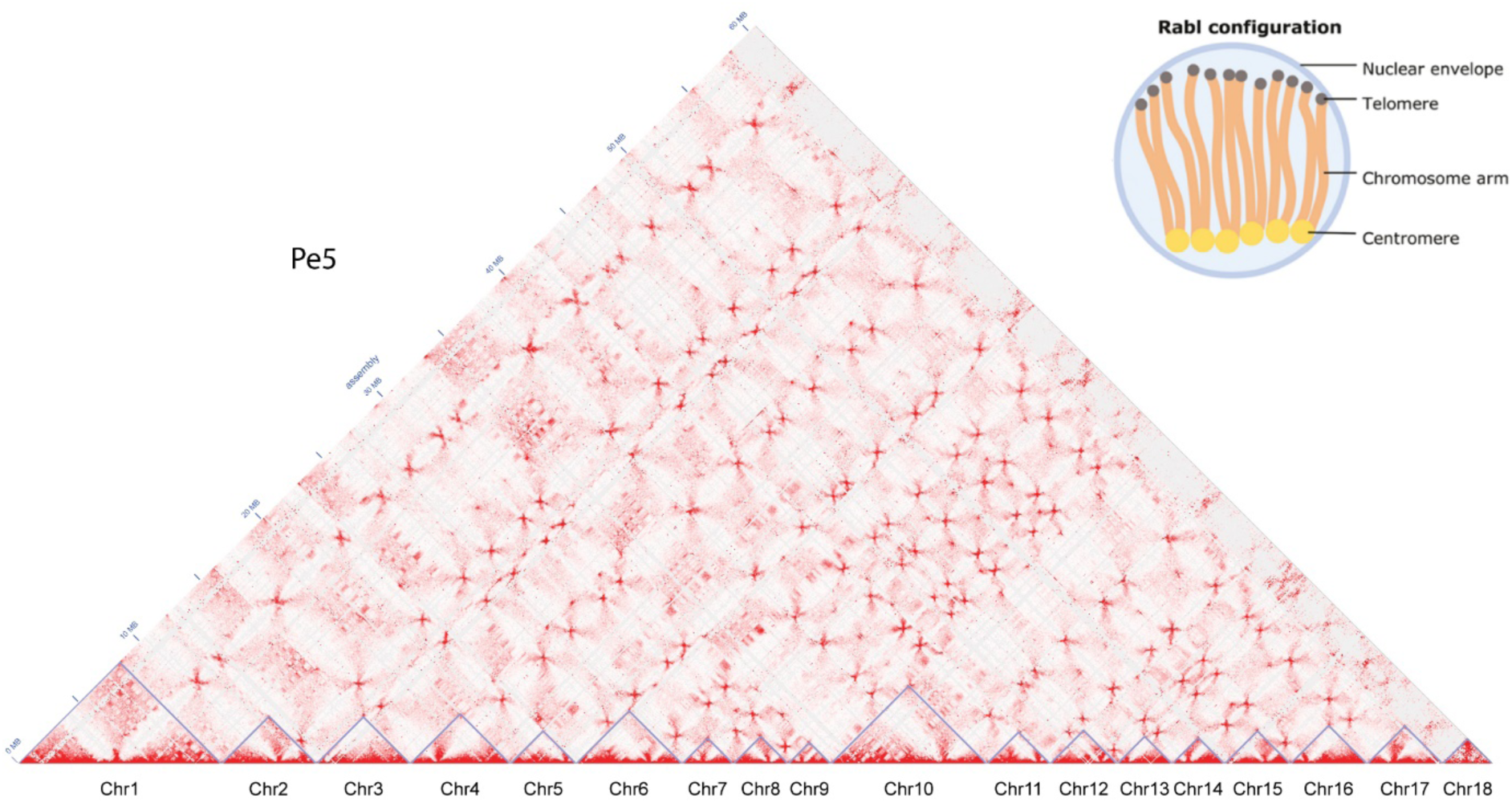
Genome-wide Hi-C contact map for the 18 chromosomes of the *Pe5* assembly. Hi-C heatmap displays the spatial interactions between chromosomes in the nucleus. Chromosome boundaries are indicated with blue lines. The observed interactions indicate a Rabl chromatin configuration with telomeric and centromeric regions inside the nucleus, as shown in the simplified schematic.

**Supplementary Figure 5.**
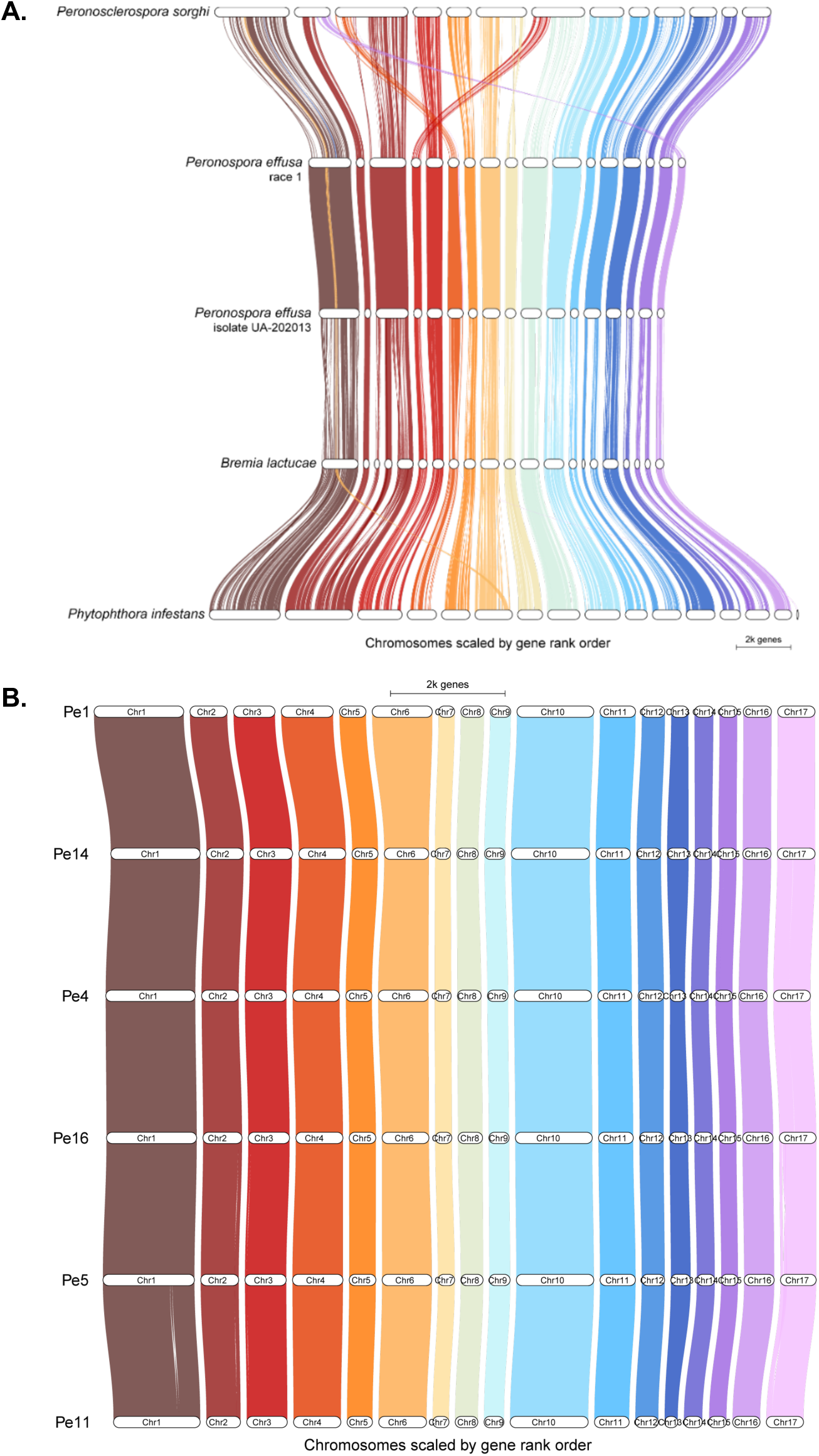
Whole-genome alignments of genomes based on sequence similarity and on relative position of protein-coding genes. **A.** Comparison of six oomycete chromosome-level genome assemblies revealing conserved chromosome structure with a few chromosomal fusions. **B.** Comparison of our six chromosome-level genome assemblies for *P. effusa* revealing highly conserved chromosome structure with two rearrangements in chromosome in *Pe11* and chromosome 17 in *Pe5*.

**Supplementary Figure 6.**
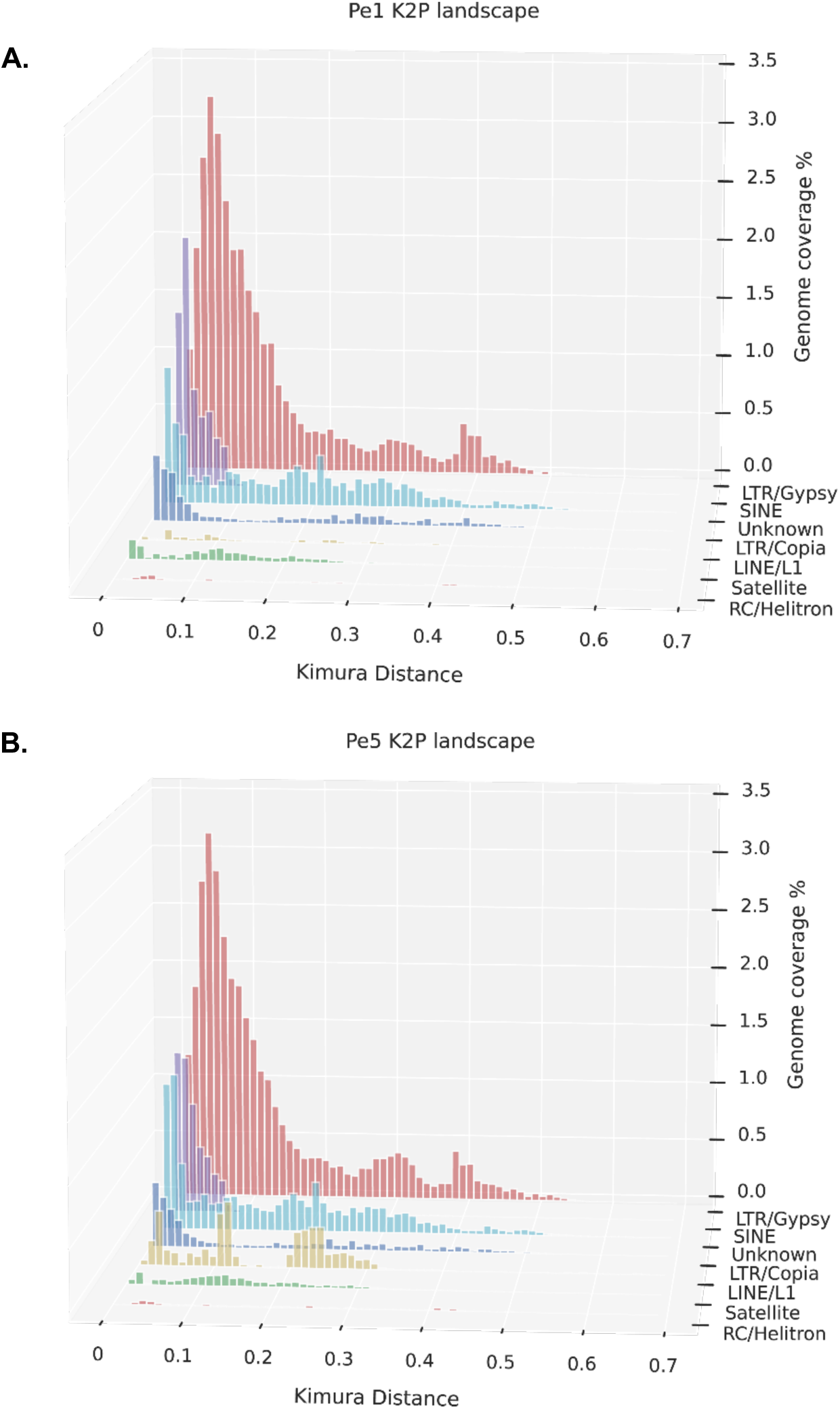
The transposable element landscape based on Kimura distances for *Pe1* and *Pe5*. Kimura distance is the measure of divergence between individual TE copies and the corresponding TE consensus sequence (Kimura, 1980), i.e., the lower the Kimura distance, the more similar the copy is to the consensus and thus the more recent it was most likely copied.

**Supplementary Figure 7.**
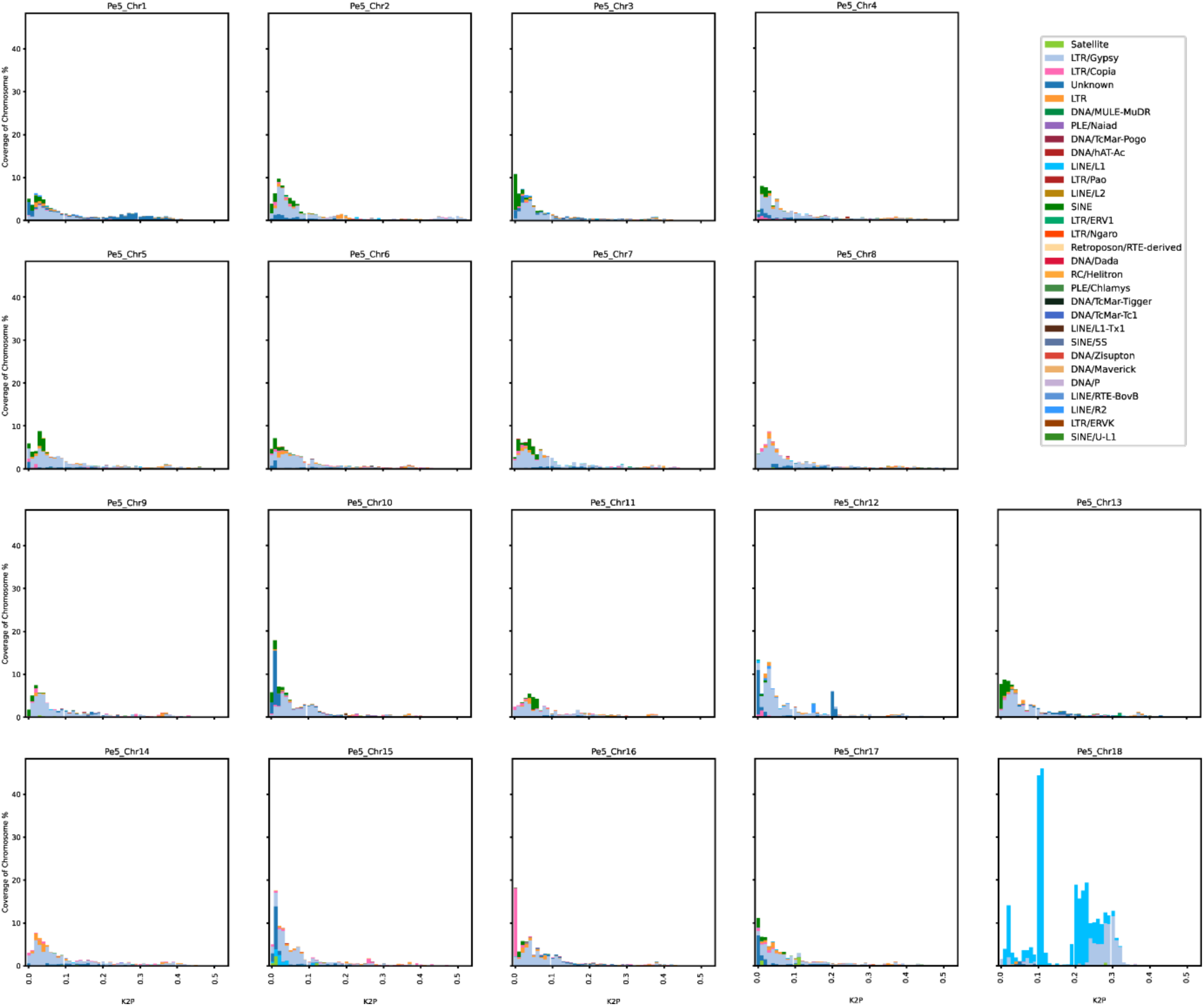
The transposable element landscape based on Kimura distances split up for each chromosome of *Pe5*. Kimura distance is the measure of divergence between individual TE copies and the corresponding TE consensus sequence (Kimura, 1980), i.e., the lower the Kimura distance, the more similar the copy is to the consensus and thus the more recent it was most likely copied.

**Supplementary Figure 8.**
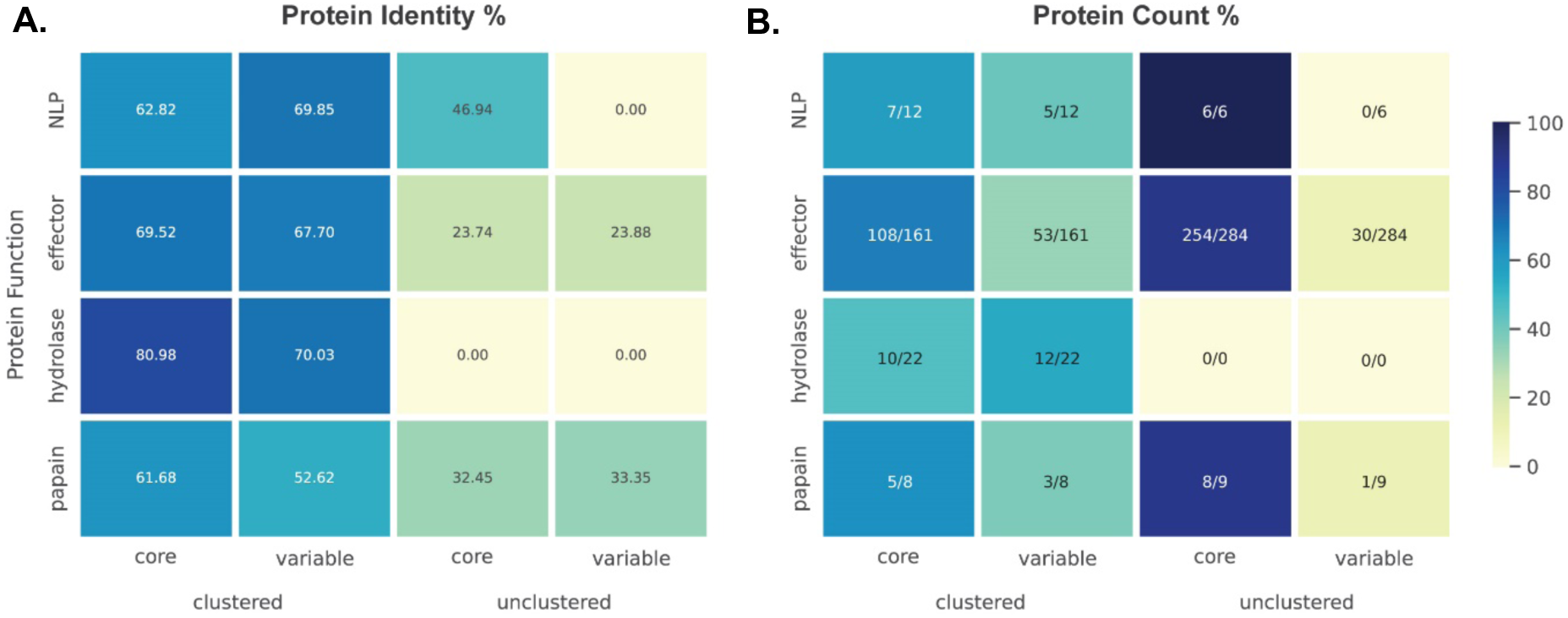
Pathogenicity-related proteins are encoded by highly variable clusters of recently copied genes. **A.**Heatmap of the average protein identity for each cluster and the average protein identity for all unclustered proteins. **B.** Heatmap of the percentage of proteins that are core or variable for each protein cluster and for all unclustered proteins.

**Supplementary Figure 9.**
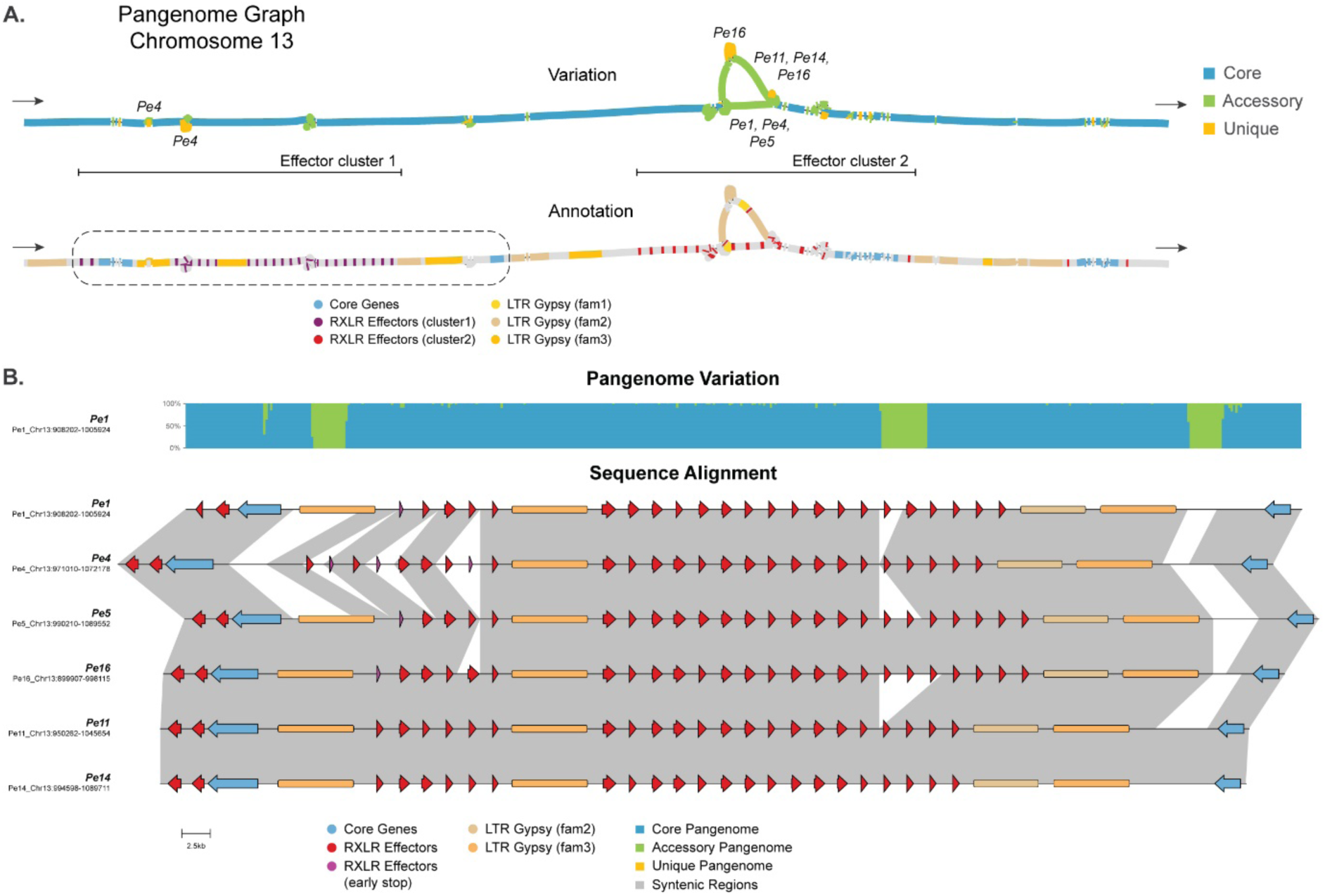
Alignment of the corresponding chromosome 13 region for six *P. effusa* isolates containing the first effector cluster. **A.** Part of chromosome 13, 0.2 Mb in size, is visualised by two pangenome graphs, with the start and end of the graph indicated with arrows. The first graph shows the variation between *P. effusa* isolates with core (blue), accessory (green), and unique (orange) regions. For large accessory and unique regions, the isolates that have these regions are indicated. The second graph visualises the effector genes belonging to the first (purple) and second (red) effector clusters on the graph. The highlighted area is expanded in **B.** Effector genes (red), pseudogenes (purple), and core genes (blue) are indicated as arrows, and LTR Gypsy repeats (fam3: orange, fam2: brown) are indicated as boxes. The pangenome variation for this region is visualised in a stacked bar plot (core blue, accessory green, and unique orange) and syntenic regions between *P. effusa* isolates are connected with grey ribbons.

**Supplementary Figure 10.**
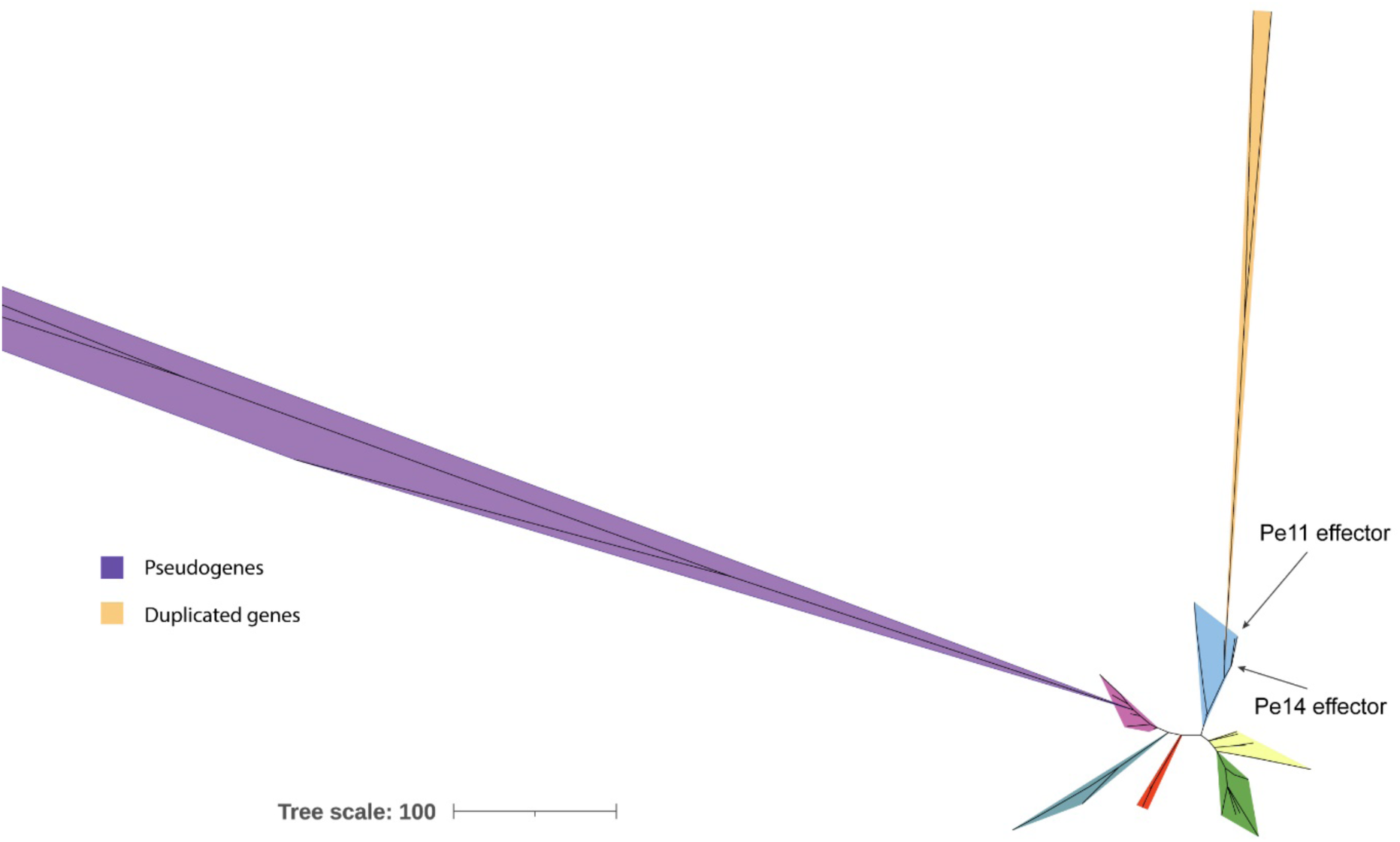
Effector gene phylogeny reveals the origin of recently duplicated genes. Nucleotide phylogeny of the 114 RXLR effector genes and five pseudogenes for the six *P. effusa* isolates in the second effector cluster on chromosome 13, highlighting the origin of the recently duplicated RXLR effectors shown in figure 7. The maximum-likelihood tree was built with IQ-TREE (v. 1.6.12) (Minh et al., 2020) and visualised with iTOL (v. 6.9) (Letunic & Bork, 2024). The location on the tree of the two genes downstream of the duplicated genes is indicated with arrows.

**Supplementary Figure 11.**
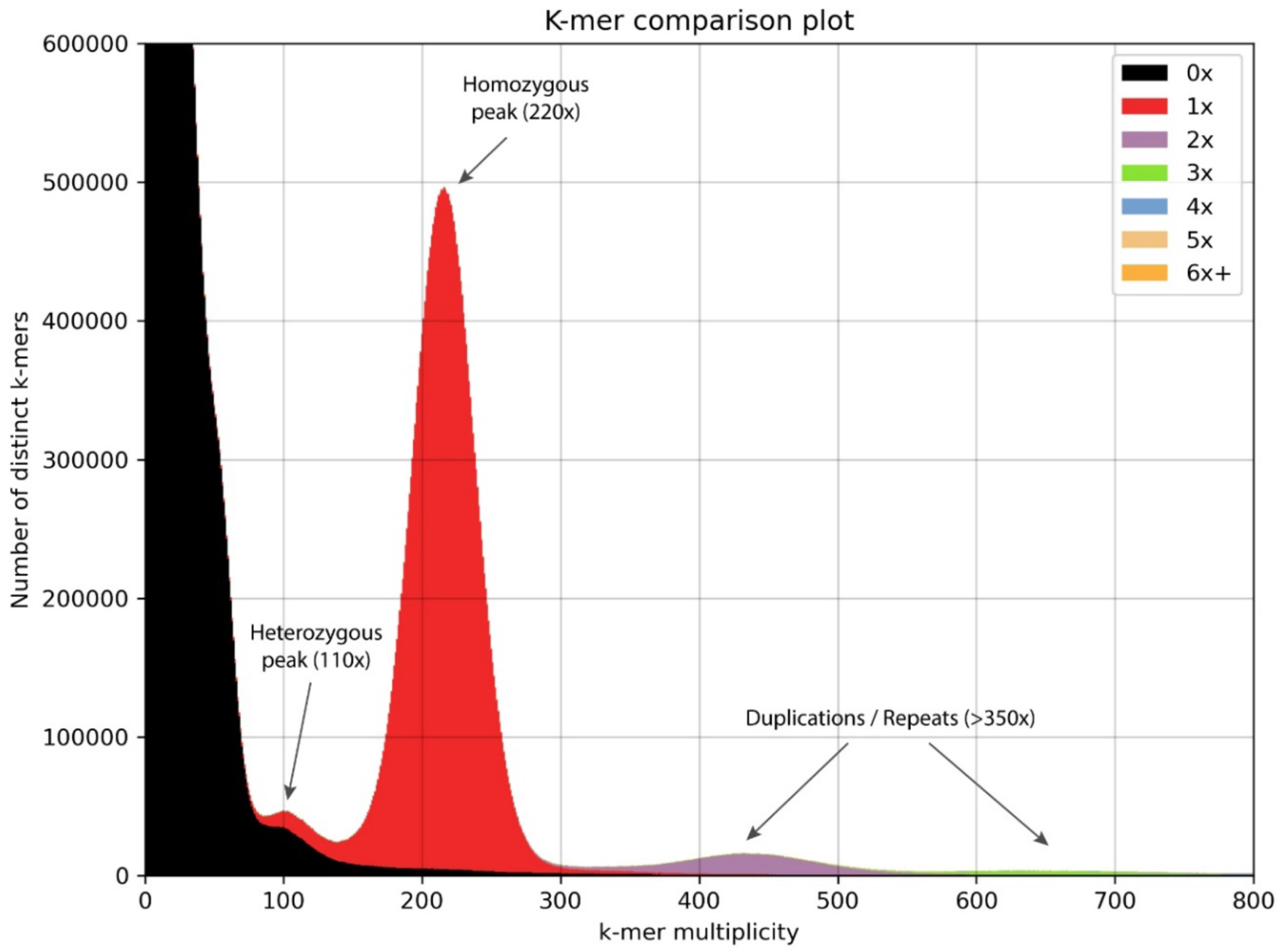
K-mer analysis confirms the completeness of the genome assembly in *Peronospora effusa* race 1 (*Pe1*). The short-read data of *Pe1* were split into K-mers (size 27b) and their frequency is plotted using KAT (v. 2.4.2) (Mapleson et al., 2017). The K-mers are mapped to the genome assembly of *Pe1* and the colour represents the number of copies that they are represented in the assembly.

**Supplementary Figure 12.**
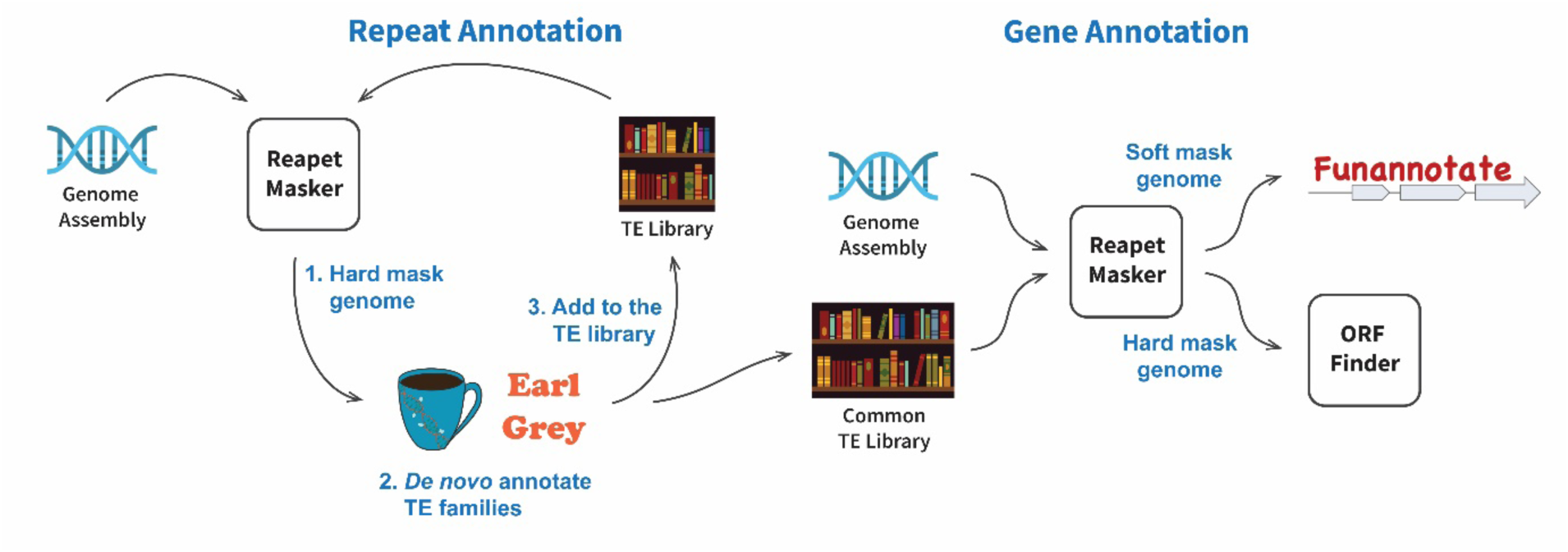
Pipeline for the separate structural genome annotation of the six *Peronospora effusa* isolates. These structural annotations were then overlapped with the pangenome graph for reference-free genome annotation and whole-genome comparisons (Figure 2).

